# Molecular architecture of the ciliary base in mammalian multiciliated cells

**DOI:** 10.64898/2026.04.05.716577

**Authors:** Caitlyn L. McCafferty, Marine Brunet, Hugo van den Hoek, Garrison Buss, Olivier Mercey, Philippe Van der Stappen, Danilo Ritz, Adrian Müller, Ricardo Righetto, Paul Guichard, Virginie Hamel, Tim Stearns, Benjamin D. Engel

## Abstract

Multiciliated epithelial cells (MCCs) generate tens to hundreds of motile cilia to drive fluid flow in diverse physiological contexts. While the axonemal structure of motile cilia has been described extensively in recent years, the molecular architecture of the transition zone, basal body, and surrounding ciliary environment of MCCs remain more elusive. Here, we use cryo-focused ion beam (cryo-FIB) milling and cryo-electron tomography (cryo-ET) to obtain *in situ* 3D views of the ciliary base within intact MCCs from mammalian trachea, complemented by *in situ* cross-linking mass spectrometry (XL/MS) and ultrastructure expansion microscopy (U-ExM) for molecular identification. Our data reveal spatially-defined modifications of microtubule architecture from the proximal centriole to the early axoneme, including transition zone-specific features such as an A-B linker bridging microtubule doublets and a helical assembly of microtubule inner proteins (MIPs). We show that the ciliary necklace, a feature observed in many motile cilia, is spatially aligned with the transition zone and quantify its regular organization within the membrane. Our *in situ* data capture rarely observed events, including intraflagellar transport (IFT) trains connecting to ciliary vesicles tethered to undocked centrioles. The surrounding ciliary environment contains intermediate filaments that encircle the basal bodies and bundled actin filaments that elaborate microvilli structures between the cilia. Integration of XL/MS and U-ExM identified novel microtubule associated proteins (MAPs), MIPs, and membrane-associated proteins localized to these distinct subdomains. This work provides a molecular and structural map of the mammalian MCC ciliary base, revealing architectural principles that underlie its assembly, organization, and function.

## Introduction

Cilia are microtubule-based organelles that project from the cell surface and mediate motility, signaling, and sensing. They fall into two broad classes: motile cilia, which beat rhythmically, and primary cilia, which are usually immotile. Motile cilia contain a complex network of proteins^1–3^ that powers coordinated beating essential for cell locomotion^4–6^, fluid transport^7,8^, and embryonic development^9,10^. In vertebrates, epithelial cells lining the trachea, brain ventricles, and reproductive tract project tens to hundreds of motile cilia on their apical surface^11^, driving mucociliary clearance^12^, cerebrospinal fluid flow^7^, and gamete transport^13^. Producing the many cilia in multiciliated epithelial cells (MCCs) involves a dedicated program of *de novo* centriole biogenesis^14^, in which centrioles form in bulk and mature into basal bodies that nucleate the formation of cilia. Defects in cilium structure and function in MCCs impair fluid movement and cause ciliopathies including primary ciliary dyskinesia^15^, chronic obstructive pulmonary disease^16^, and hydrocephalus^17^.

Cilia are organized into two principal regions: the basal body and the axoneme. The basal body, derived from the mother centriole, templates the cilium and docks it to the plasma membrane via distal appendages^18^. The axoneme is composed of microtubules and associated proteins and extends from the basal body. In motile cilia, the axoneme has additional structures, such as a central pair, radial spokes and dynein arms^3^ that are involved in generating the ciliary stroke. At the base of the axoneme, just distal to the basal body, the transition zone forms a selective gate that regulates molecular traffic between the cytoplasm and the ciliary compartment^19^.

Electron microscopy (EM) has long been central to studying multiciliated tissue *in situ*^20–25^. The more recent application of EM to image vitreously frozen samples in three dimensions by cryo-electron tomography (cryo-ET)^26,27^ has revealed molecular details of isolated basal bodies and transition zones from MCCs of *T. thermophila*, Chinese hamster ovary, bovine trachea epithelia, and mouse photoreceptor cilia^28–32^. Thinning frozen samples with a focused ion beam (FIB)^33,34^ has enabled cryo-ET of the ciliary base inside cells (i.e., *in situ*), including in *C. reinhardtii*^35^, mammalian sperm^36^, and mouse ependymal MCCs^37^, providing native views of complex biological processes such as the mass production of centrioles from deuterosomes^37^ and the stepwise assembly of intraflagellar transport (IFT) trains before entry into the cilium^35^. Although these studies have provided important structural insights into the components of the ciliary base, we still lack an integrated map of this cellular region in MCCs.

In addition to visualizing native cellular landscapes, *in situ* cryo-ET data can be used to resolve molecular structures in cells by subtomogram averaging (STA). However, FIB milling and cryo-ET have relatively low throughput for rare and challenging targets such as the ciliary base, making it difficult to attain STA resolution that is high enough for *de novo* protein identification. This limitation can be addressed by integrating cryo-ET with complementary approaches for investigating molecular identification, structure, and localization *in situ*, including cross-linking mass spectrometry (XL/MS) and ultrastructure expansion microscopy (U-ExM)^38^. XL/MS provides amino acid-level mapping of protein interactions and can be performed *in situ* using commercially available, membrane-permeable chemical cross-linkers^3,39,40^. U-ExM improves epitope accessibility and resolution of immuno-stained proteins *in situ* by physically expanding the sample^41,42^, allowing the molecular identification of specific substructures when correlated with cryo-ET^35,43^.

Here, we apply an integrative *in situ* approach^38^ to the ciliary base of mammalian MCCs. We use FIB milling and cryo-ET to explore the native molecular architecture of multiciliated mouse trachea epithelial cells (mTECs), a well-studied type of MCC. We complement our cryo-ET data with *in situ* XL/MS and U-ExM, generating an extensive protein interaction network of mammalian MCCs. These integrated methods reveal structural changes of microtubules and associated proteins in a spatially-defined pattern from basal body to transition zone and axoneme, the specific localization of several proteins associated with this ciliary ultrastructure, and the precise organization of the ciliary necklace, a defining ultrastructural feature of the ciliary membrane. Moreover, we uncover IFT train interactions at the ciliary base and characterize the arrangement of cytoskeletal filaments, particularly actin bundles, surrounding the basal bodies and forming microvilli structures between adjacent cilia. Together, our data provide an integrated map of the ciliary base in MCCs and provide a rich resource of validated protein-protein interactions.

## Results

### Cryo-ET of MCCs reveals the native architecture of the ciliary base

To visualize the ciliary base *in situ*, we performed cryo-ET on FIB-milled mTECs grown and differentiated in culture (**Figures S1 & S2**). This approach allowed us to acquire tomograms of the apical surface and cellular interior containing multiple cilia, which we analyzed by membrane segmentation, filament tracing, STA, classification, and tomogram map-backs of several different ciliary base protein complexes (**Figure 1A,F**). We observed IFT trains, presumed to be anterograde based on their extended morphology, assembling at the ciliary base and along the axonemal microtubules (**Figure 1B**). Bundled actin filaments were found in close proximity to the ciliary membrane (**Figure 1C**). We also observed thicker intermediate filaments (IFs) forming a network around the proximal region of the centrioles (**Figure 1E**). Embedded in the membrane at the base of the cilium, we identified periodic membrane-associated densities consistent with the ciliary necklace complex (**Figure 1D**).

**Figure 1:**
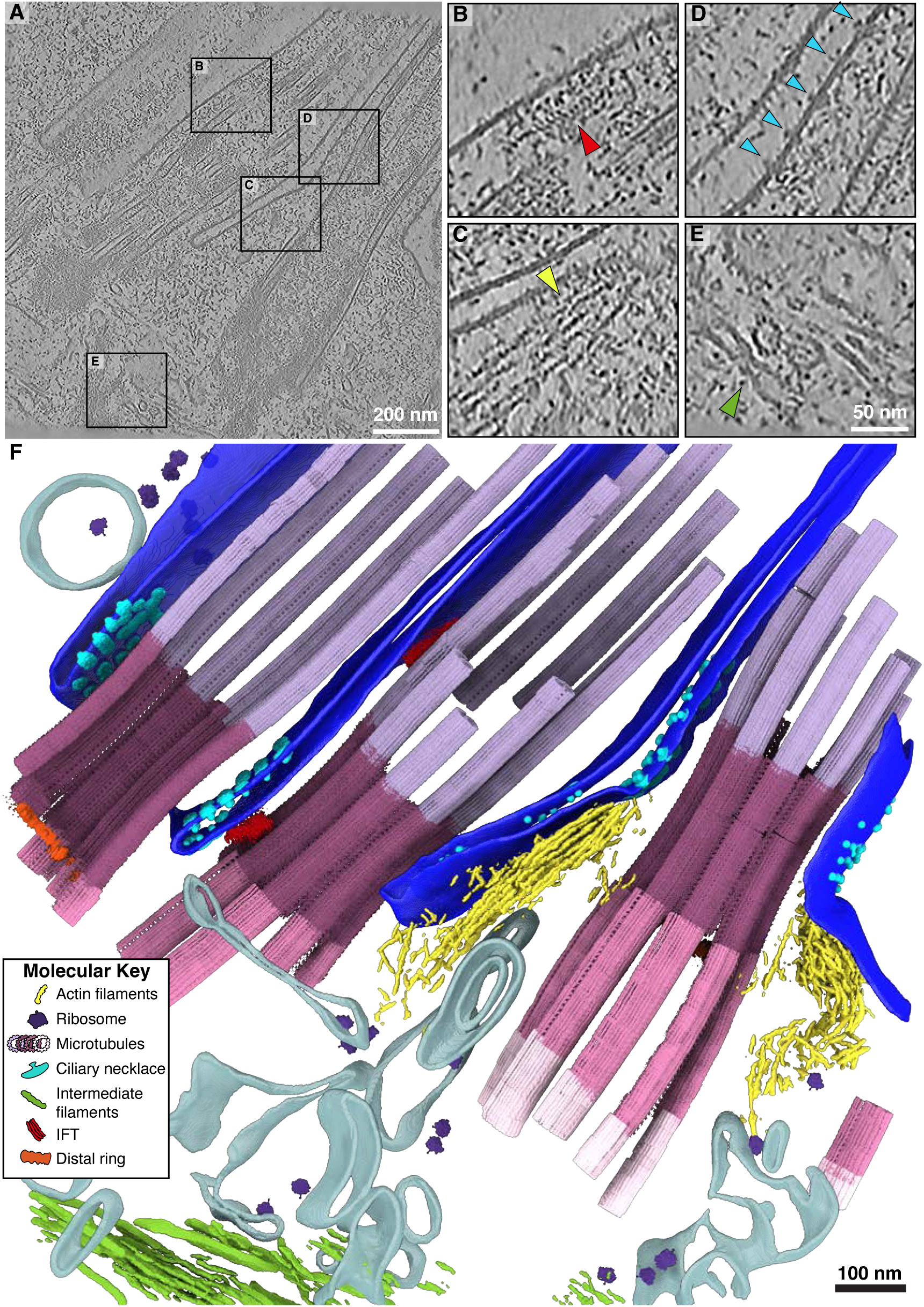
Cryo-ET of multiciliated mouse tracheal cells (mTEC) provides an atlas of cellular structural biology of the ciliary base. **A)** Slice through a representative tomogram from our mTEC dataset, showing the multiciliated apical surface of a mammalian airway epithelial cell. **B)** A zoom in on an intraflagellar transport (IFT) train traveling along the microtubules of the cilia. **C)** Actin filaments positioned beneath the apical membrane, interacting with the ciliary pocket. **D)** Membrane densities belonging to the ciliary necklace localize in the ciliary pocket, adjacent to the transition zone. **E)** Intermediate filaments are observed at the basal body level (below actin) under the centriolar base. **F)** Segmentation and map back of subtomogram averages of the tomogram in **(A)**. The combined rendering displays the rich molecular contents of the dataset including actin filaments, ribosomes, distinct microtubule classes (shades of pink and purple as in Fig. 2), ciliary necklace, intermediate filaments, intraflagellar transport (IFT) trains, and the centriolar distal ring. The ciliary membrane is shown in royal blue while other membranes and vesicles are tan.

To map ultrastructural changes from the basal body to the proximal portion of the axoneme, we traced ciliary microtubules along their entire length, then performed STA and classification (**Figures 2A-E**). Five discrete structural classes emerged, each corresponding to a specific subregion of the cilium. A sliding-window reassignment approach sharpened these boundaries, yielding four well-defined regions—proximal basal body, core/distal basal body, transition zone, and early axoneme—plus a short boundary class between the distal basal body and transition zone, coinciding with the switch from triplet to doublet microtubules and the position of the basal body’s distal luminal ring^43^ (**Figure 2F**). Diameter measurements showed that the nine-microtubule barrel narrows from the basal body to the transition zone before expanding again at the early axoneme (**Figure 2G**).

**Figure 2:**
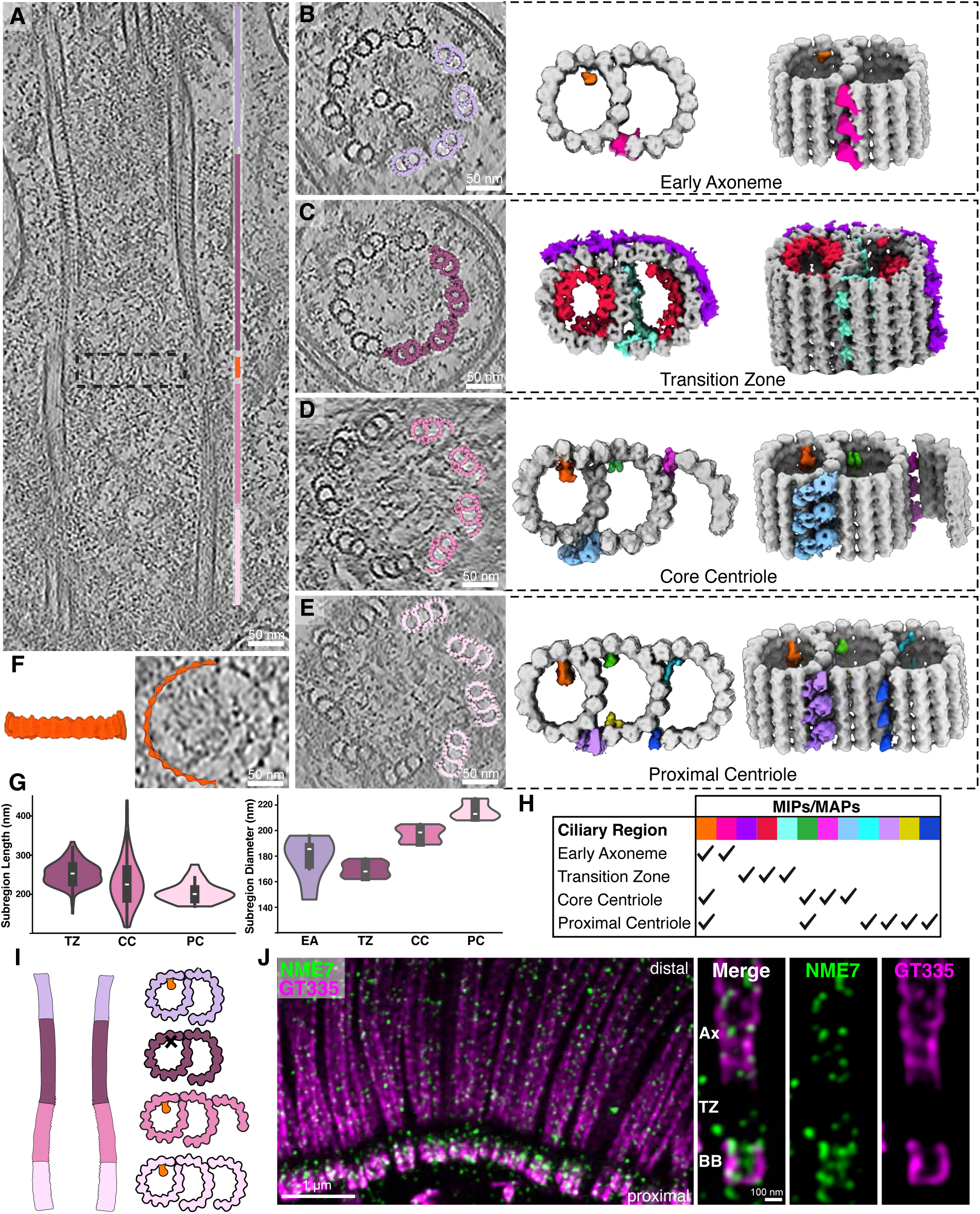
Zonal specialization of the ciliary base revealed by structural transitions along the length of the basal body and cilium. **A)** Cryo-ET longitudinal slice through a cilium from mTEC showing distinct subregions: proximal centriole (light pink), core centriole (pink), hook/distal ring region (orange), transition zone (plum), early axoneme (lilac). **B-E)** Cross-section slices of regions shown in **(A)**, illustrating the compositional changes in the microtubule ultrastructure from triplet to doublet and the appearance/disappearance of specialized microtubule associated densities. **F)** Average of the distal ring, which sits between the transition zone and the distal centriole^43^. **G)** Length distributions of the proximal centriole (mean: 205 nm), core centriole (mean: 230 nm), and transition zone (mean: 253 nm) subregions from tomograms (left). Distributions of diameters for the proximal centriole (mean: 215 nm), core centriole (mean: 197 nm), transition zone (mean: 169 nm), and early axoneme (mean: 178 nm) subregions (right). The length of the transition zone measured by U-ExM (mean: 274 nm) is consistent with our cryo-ET measurements. **H)** Table summarizing the presence of different MIPs/MAPs shown in (**B**)-(**E**). Orange: NME7; Red: MIP helix. **I)** Diagram of A-tubule MIP in subtomogram averages along the microtubules. **J)** U-ExM of human multiciliated tracheal cells (hTEC) stained for NME7 (green) and polyglutamylated tubulin (GT335, magenta) shows that NME7 localizes to the centriole and axoneme, but not the transition zone.

Each subregion displayed unique architectural signatures. Moving from proximal to distal, we observed progressive loss of the C tubule (**Figure 2B-E**) and variations in the decorating MIPs and MAPs (**Figure 2H**). The basal body showed the canonical A–C linker (**Figure S3A-C**)^29,32,44^ and a MAP between the A- and B-tubules that closely resembles the “stem” portion of the inner scaffold that has been observed in several other species^45^. In addition, we identified a preserved A-tubule MIP at position A9 that is also present in the core centriole and axoneme with an 8 nm repeat, but notably absent in the transition zone (**Figure 2I**). The position of the density in the A-tubule suggests that it corresponds to NME7^46^. Consistent with this hypothesis, U-ExM revealed that NME7 in human tracheal epithelial cells (hTECs) was present at both the basal body and axoneme but absent from the transition zone, matching the pattern we saw by cryo-ET (**Figures 2J & S4)**. Furthermore, docking an AlphaFold-predicted NME7 structure into the proximal basal body density (**Figure S3D**) supported this assignment, in agreement with several independent studies^37^. To further test NME7 localization, we stained for the protein in hTERT RPE-1 cells using U-ExM and found that it localizes to the centriole exclusively, confirming this protein as both a motile and primary cilia centriolar MIP (**Figure S5**).

### An interactome of mammalian MCC cilia uncovers microtubule-, actin-, and membrane-associated networks

While cryo-ET and U-ExM allowed us to map the structural transitions of microtubules across the ciliary base, these methods alone did not reveal new ciliary protein identities. In addition, because it relies on averaging many particles, STA can fail to resolve density for proteins with asymmetric radial distributions or sub-stoichiometric binding along the microtubules. To address this, we turned to in situ XL/MS, which provides complementary information on protein identity and interaction networks.

Our *in situ* MCC interactome was generated by adding a membrane permeable, MS cleavable cross-linker to detached cilia and basal bodies. To obtain sufficient material for *in situ* XL/MS, we isolated basal bodies and cilia from bovine trachea (bTEC) (**Figure S1**). The resulting dataset comprised more than 10,500 unique cross-links within 1,661 proteins, representing 1,171 protein-protein interactions (**Figure 3A & Table S1**). To enrich for distinct subregions of the cilium, we employed four isolation strategies, collecting fractions enriched for: (1) whole cilia including basal bodies (BBC), (2) basal bodies alone (BB), (3) membrane and matrix (MM), and (4) microtubules (MT; described in **Methods**). Gene ontology (GO) analysis of these fractions confirmed selective recovery of the intended compartments (**Figure 3B**), providing the first XL/MS–based interactome of mammalian motile cilia, including the basal body, transition zone, cytoskeletal, and membrane-associated protein networks.

**Figure 3:**
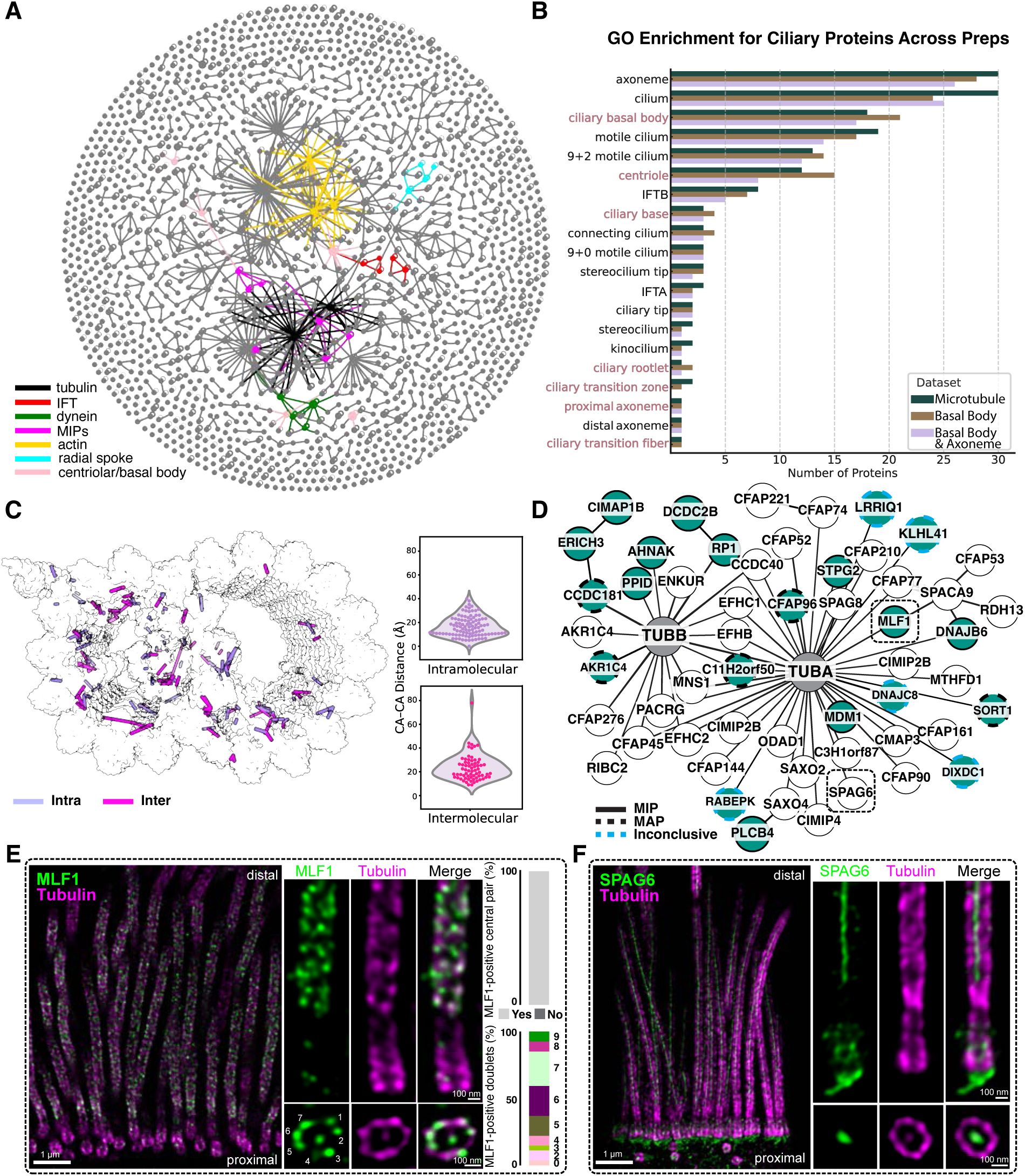
Mapping the tracheal cilia interactome using cross-linking mass spectrometry (XL/MS) and expansion microscopy (U-ExM). **A)** XL/MS of bovine tracheal epithelial cells (bTEC) reveals an interactome of over 1,600 proteins with over 10,500 unique amino acid links. The interactome was generated using three biochemical enrichment strategies targeting the basal body and cilia (BBC), basal body (BB), or the microtubules (MT) (**Methods**). We highlight a few proteins belonging to different ciliary complexes (this is not exhaustive). **B)** Gene ontology (GO) enrichment analysis of proteins from each condition. BB-enriched fractions indeed show an enrichment for basal body/centriole proteins (highlighted in dusty pink). **C)** As an internal control, intramolecular and intermolecular XLs that are represented on isolated DMTs from bTEC are measured and displayed on the structure. **D)** Proteins that we found XLed to tubulin in our interaction network. Several tubulin binding proteins that are not present in the structure in **(C)** are shown in turquoise. XL positions can be used to map the binding proteins to the interior or exterior of the microtubule. The mapping of the turquoise tubulin binding proteins is shown by the node outlines. **E)** U-ExM localizes MLF1, a previously uncharacterized MIP, to the axoneme of hTEC. Side views (top): of MLF1 staining (green) co-localized with axoneme microtubules (tubulin: magenta). The cross-section views (bottom): MLF1 always binds to the central apparatus and most often 6-7 of the 9 DMTs (**Fig S6**). **F)** U-ExM of SPAG6 (green) shows that the protein is not only localized to the central apparatus, but also the ciliary base near the basal foot/rootlets^56^.

We next validated the dataset by mapping cross-links onto a published single particle analysis (SPA) structure of isolated doublet microtubules (DMTs) from bovine trachea^47^. 94% of the 169 mapped cross-links fell within the 35 Å distance restraint of the linker (**Figure 3C**), emphasizing the structural specificity of the data. Within the interactome, we identified 46 proteins cross-linked to tubulin, including several not previously annotated as tubulin-binding proteins (**Figure 3D**). To confirm the *in situ* localization, we used U-ExM to label candidate MIPs and MAPs (**Figure S6-S9**).

Among the proteins in our network, MLF1 stood out for its extensive cross-links to both alpha tubulin (TUBA) and microtubule-binding proteins. MLF1 belongs to a poorly characterized protein family associated with immune response, apoptosis, and cell cycle regulation^48–50^, and it has been implicated as both a tumor suppressor and an oncogene^48^. Although primarily studied in the context of cancer, MLF1 has been localized to primary cilia and mTECs by immunofluorescence^51^, and to photoreceptor cilia by proteomics^52^. Furthermore, transcriptomics^51^ shows it is upregulated during ciliogenesis, and it has been proposed as a candidate ciliopathy gene^53^. Here, XL/MS identified a cross-link to TUBA (position 370), indicative of binding within the microtubule lumen. U-ExM confirmed MLF1 localization to axonemal microtubules and the central apparatus (**Figures 3E & S6**). While it was consistently present in the central apparatus, MLF1 was variably associated with the nine microtubule doublets. The occupancy changed along the length of the cilium, suggesting that sub-stoichiometric association may be why the protein had not been identified in previous axonemal structures (**Figure 3E**). The previous localization of this protein to primary cilia prompted us to stain for the protein in hTERT RPE-1 cells by U-ExM, however we observed no clear localization in the primary cilia of these cells (**Figure S5**). Together, these data position MLF1 as a previously unrecognized MIP in mammalian MCCs and suggest a specialized role in motile cilia, likely spatially restricted within the axoneme.

We also investigated the localization of SPAG6, an armadillo repeat protein, linked to TUBA and CFAP69 in our interactome (**Figures 3D, S7 & Table S1**). SPAG6 has been localized to the central apparatus in *C. reinhardtii*^54^ and to both ciliated and non-ciliated cells in mouse tissues, suggesting a role independent of the central apparatus^55^. Indeed, studies of *Spag6*-deficient mice showed fewer cilia per tracheal cell and a reduction in cilia beat frequency, suggesting that this protein functions in motility through an association with the central apparatus, but also has a role in ciliogenesis and cell polarity^56^. Our U-ExM analysis of SPAG6 in hTEC confirmed localization of the protein to the central apparatus, and unexpectedly, to the basal body and its sub-proximal region, consistent with rootlet association (**Figure 3F**). As anticipated, the protein was not localized to primary cilia in hTERT RPE-1 cells (**Figure S5**).

Finally, our dataset contains an extensive set of MAPs, including CFAP58 and CCDC146, which we previously identified^3^. These proteins were cross-linked to additional ciliary proteins predicted to bind the microtubule exterior. We identified RP1 as a microtubule associated protein that is present in respiratory MCCs but absent in hTERT RPE-1 (**Figure S5 & Table S1**). We also detected cross-links between calmodulin and multiple axonemal and membrane-associated proteins, consistent with its proposed regulatory role in MCC function^57^ (**Table S1)**. Together, these analyses define a molecular interactome of mammalian MCCs, uncovering links between cytoskeletal, membrane, and regulatory networks.

### The A-B linker and MIP helix define a distinct architecture of the mTEC transition zone

The transition zone marks a major structural shift at the base of the cilium. We observed that termination of the C-tubule of the basal body coincides precisely with the position of the luminal distal ring and the emergence of an A–B linker bridging adjacent microtubule doublets (**Figure 4A,C**, pink), as the diameter of the transition zone narrows relative to the bordering basal body and axoneme (**Figure 2G**). Transition zone microtubules show a loss of the A-tubule MIP that we identified as NME7 (**Figure 2H-J**) and the appearance of a helical MIP decoration inside both A- and B-tubules, which we call the MIP helix^31^ (**Figure 4A,B**, red). Additional MAPs decorate the exterior of the A- and B-tubules of the transition zone, including a helical wrap around part of the B-tubule and additional density around the A-tubule (**Figure 4A,D**, purple). In the transition to the early axoneme, the A–B linker and MIP helix are lost, and the NME7 A-tubule MIP reappears. Notably, the transition zone shows only minimal polyglutamylated tubulin (GT335) signal, distinguishing it from the basal body and axoneme, as previously described in *Chlamydomonas*^35^ (**Figures 2 & S4**).

**Figure 4:**
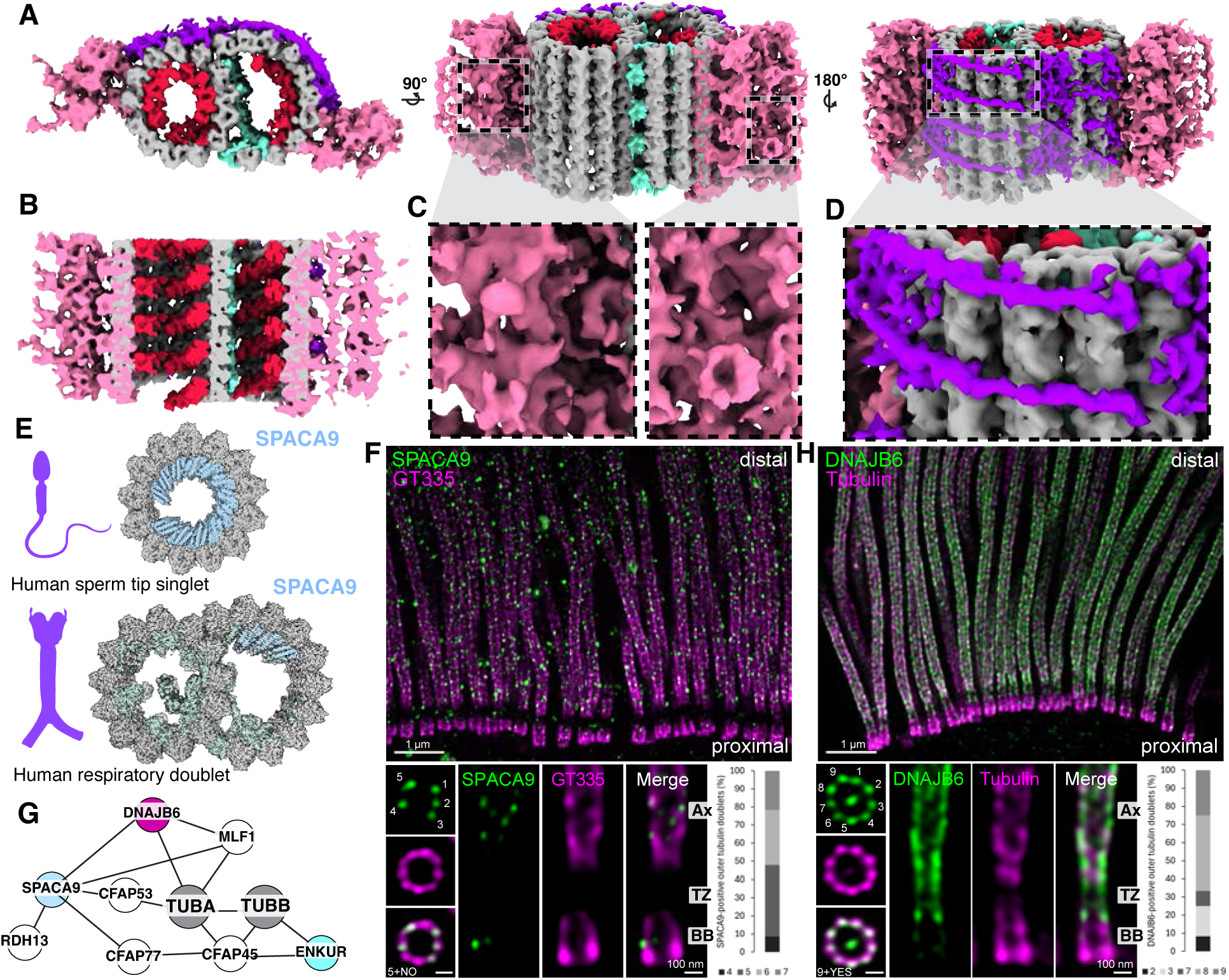
The transition zone contains a distinct A-B linker and MIP helix. **A)** Subtomogram average of the mTEC transition zone combining STA maps centered on the microtubules (16 Å resolution) and linker (24 Å resolution), with tubulin densities in grey, the A-B linker in pink, outer helical wrap in purple, and inner helical protein (MIP helix) in red. **B)** Clipped view through the STA map showing the MIP helix that binds both the A and B tubule of the transition zone. **C)** Inset of the A-B linker, showing different views of the density. **D)** Inset of the outer helical wrap protein. **E)** Single-particle analysis structures from other studies show the binding of SPACA9 in sperm and respiratory axonemes^58^. **F)** U-ExM of hTEC cells stained with SPACA9 (green) and polyglutamylated tubulin (GT335, magenta) showing the localization of SPACA9 at the axoneme. SPACA9 binds 4 to 7 of the 9 DMTs, based on top-view images (n = 23). **G)** XL/MS network from bTEC highlighting a subnetwork from Figure 3, including SPACA9, a protein that has a similar topology to the inner helical density. **H)** U-ExM of hTEC cells stained with DNAJB6 (green) and tubulin (magenta). DNAJB6 localizes to both the transition zone and axoneme. Quantification of DNAJB6 foci associated with the 9 DMTs. The most frequent category corresponds to eight foci, representing ∼42% of the cases (n = 12). Basal body (BB), transition zone (TZ), axoneme (Ax).

The MIP helix density in our STA resembles the SPACA9 density previously described in human sperm tail tips and respiratory doublet microtubules^58–60^ (**Figure 4E**). Interestingly, U-ExM revealed that SPACA9 localizes to the axoneme (consistent with a previous SPA structure^58^), although lack of staining in primary cilia suggests that SPACA9 is a motile cilia-specific MIP (**Figures 4F & S5**). However, U-ExM showed no enrichment of SPACA9 at the hTEC transition zone, suggesting that this protein is an unlikely candidate for the MIP helix (**Figures 4F & S8**). We next examined other candidates from our XL/MS network, focusing on DNAJB6, a tubulin-binding protein extensively linked to SPACA9 and MLF1 in our interactome (**Figure 4G**). Unlike SPACA9 and MLF1, DNAJB6 localized to both the transition zone and axoneme in hTECs (**Figures 4H & S9**). The most prevalent pattern involves decoration of 8 doublets, accounting for 42% of the observed top views. Additionally, DNAJB6 shows robust localization at the central apparatus of the axoneme (**Figure S9**). Previous AP/MS found that DNAJB6 interacts with transition zone protein RPGRIP1L^56^, suggesting its significance in the transition zone; whereas the cross-links between DNAJB6 and both SPACA9 and MLF1 indicate possible interactions between these proteins in the axoneme. Staining for DNAJB6 in primary cilia of hTERT RPE-1 cells showed the opposite localization relative to motile cilia, with the protein only present at the basal body and absent from the transition zone and axoneme (**Figure S5**).

Together, these results reveal that the mTEC transition zone is defined by a combination of structural features—an A–B linker, a MIP helix, distinct MAP decorations including a helical wrap, and lack of glutamylation—that are absent from adjacent regions of the cilium. Proteomic and localization data allowed us to pinpoint DNAJB6 as a likely transition zone-associated microtubule-binding protein with a potential stabilizing or regulatory role.

### The ciliary necklace forms a regularly spaced decoration around the transition zone

Another striking observation from our cryo-ET data was the presence of a regularly spaced membrane protein complex adjacent to the transition zone (**Figure 5A-C**), consistent with the “ciliary necklace” first visualized in motile cilia by freeze-fracture and freeze-etching^61,62^. Similar structures have also been described in the connecting cilium of photoreceptors^32^.

**Figure 5:**
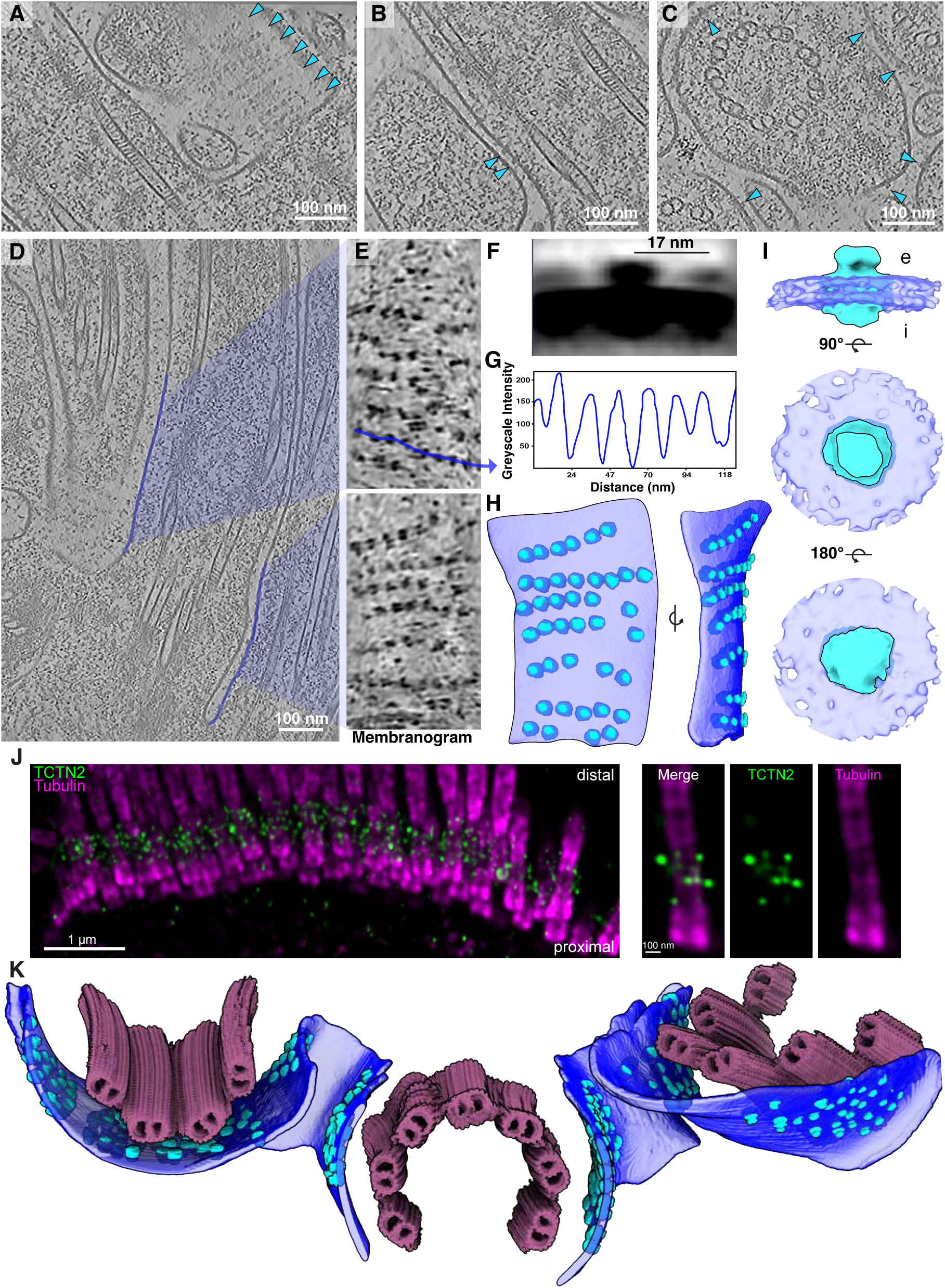
Subtomogram averaging of the mTEC ciliary necklace highlights periodic membrane complexes and potential coordination with the transition zone. **A-D)** Cryo-ET slices showing views of ciliary necklace particles (light blue arrowheads): **(A)** top view slice**, (B)** slice along the longitudinal axis, **(C)** cross-section slice. **D)** Segmented membranes are highlighted with blue lines and were used to create **E)** the flattened ciliary membrane (membranogram)^63,112^ displaying distinct particles of the ciliary necklace organized into 6-7 rows. **F)** Central slice through the ciliary necklace STA map, showing 17 nm between adjacent particles. **G)** Line profile across the flattened membrane (from blue line in E), showing distances between particles in a single row, consistent with the 17 nm determined by STA. **H)** Segmented membrane and particle mapbacks corresponding to the lower membranogram in **(E)**. **I)** Isosurface renderings of the STA map of an individual ciliary necklace unit, revealing the membrane-spanning complex with protein densities extending into the intracellular (i) and extracellular (e) spaces. **J**) U-ExM of hTEC cells stained with TCTN2 (green) and tubulin (magenta) highlighting its restricted localization at the transition zone. **K)** Ciliary necklace and transition zone mapbacks show the spatial alignment between the two structures, suggesting that the ciliary necklace may tether the transition zone to the ciliary membrane.

To better visualize the putative ciliary necklace, we used “membranograms”^63^ to look at the flattened surface of the ciliary membrane. This revealed distinct puncta of protruding densities arranged in concentric rows on the membrane surface (**Figure 5D,E**). Manual picking and quantification showed an average of 7 ± 0.68 rows per cilium (n=15), consistent with previous freeze-fracture measurements of cilia in rat tracheal cells that identified 6 rows of the necklace^61^. Line-profile analysis across the flattened membrane indicated a mean center-to-center spacing of 17.50 ± 0.30 nm (n=218) between adjacent necklace particles (**Figure 5G,H**), consistent with the spacing measured directly from STA of these particles (∼17 nm) (**Figure 5F**). This spacing of 17 nm suggests that there is a stoichiometry of approximately 6 necklace particles to each doublet microtubule.

STA revealed each necklace particle to have a ∼6 nm extracellular density with a broader ∼9 nm domain partially embedded within the membrane and extending only slightly into the transition zone matrix (**Figure 5I**). Although the resolution of the average was insufficient for unambiguous molecular identification, prior work has localized proteins such as CEP290^64^, TMEM67^65^, and TCTN2^65^ to this region. Our U-ExM data clearly shows TCTN2 localized at the transition zone in a punctate pattern, however, the resolution of the immuno-staining does not allow for reliable measurement of center-to-center spacing between individual particles (**Figure 5J**).

The ciliary necklace has been proposed to physically interact with the Y-links of the transition zone. However, in our mTEC tomograms, Y-link densities are not visible likely due to their flexibility, and focused STA revealed only small extensions beyond the doublet microtubules (**Figure S3E**). Side views of transition zones in the tomograms lacked the ordered Y-link architecture described in conventional TEM of photoreceptor cilia, precluding direct visualization of a necklace–Y-link connection. Despite not being able to average the Y-links, mapping of our ciliary necklace particles and transition zone particles onto the tomograms demonstrated a spatial coordination between the necklace and the transition zone microtubules (**Figure 5K**).

Our structural and proteomic analysis of the ciliary necklace defines its precise arrangement around the transition zone and suggests potential molecular connections to the underlying Y-link architecture. These findings highlight the complex supramolecular organization of the membrane–microtubule interface at the ciliary base, where gating and trafficking machinery converge.

### IFT trains interact with ciliary vesicles and assemble at the base of mature cilia

Our cryo-ET of mTECs revealed what appeared to be assembling IFT trains associated with transition zone microtubules at the base of several mature cilia, similar to observations from *C. reinhardtii*^35^ (**Figures 6A & S10A-B**). Indeed, STA of these IFT trains, although limited in resolution due to low particle number, revealed features that match the highly conserved anterograde train structure previously resolved in *C. reinhardtii*^66^. Consistent with this cryo-ET observation, cross-links that we found within the IFT proteins from the isolated cilia show correspondence with a published anterograde IFT structure^67^ (**Figure 6B,C**). We also identified an intermolecular cross-link between Ezrin and IFT88, a core component of the IFT-B complex (**Figure 6D**). Ezrin sits at the base of the ciliary membrane in MCCs, anchoring filamentous actin to the membrane^68^. This suggests a physical link between the ciliary membrane and IFT assembly machinery.

**Figure 6:**
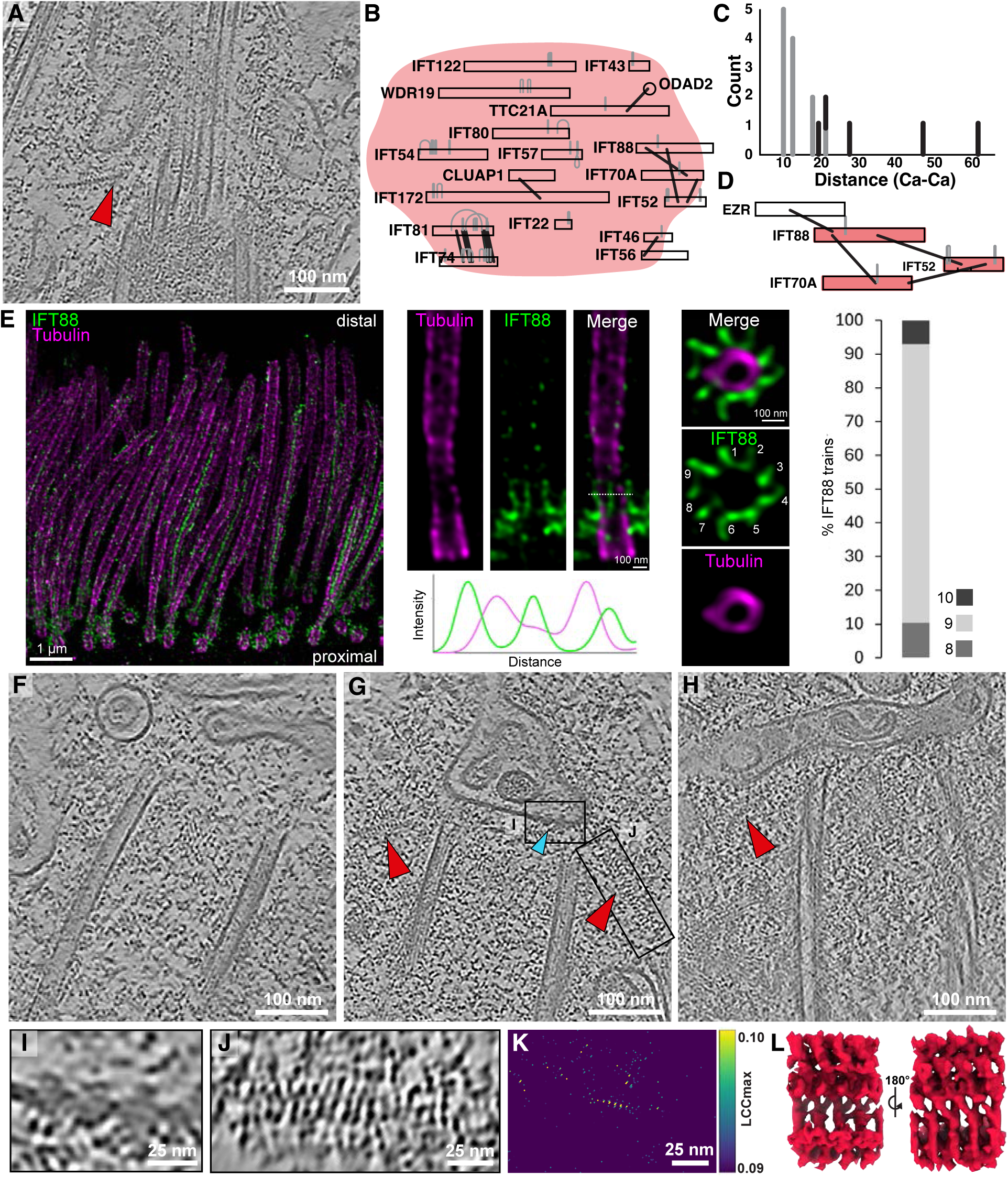
Visualization of undocked centrioles with vesicle-associated IFT trains and molecular interactions suggest pre-ciliary assembly coordination. **A)** Cryo-ET slice showing an IFT train (red arrowhead) at the base of a mature cilium. **B)** XLs of IFT-A and IFT-B proteins from bTEC cilia. **C)** Distances between XL pairs measured on the IFT-B structure from *C. reinhardtii* reinforce the conserved structure between eukaryotic clades. **D)** An intermolecular XL connects IFT88 to Ezrin, a known membrane-cytoskeletal linker. **E)** U-ExM localizes IFT88 (green) to the base of mature hTEC cilia (tubulin: magenta), consistent with our observations in cryo-ET. Quantification of the number of IFT88 trains attached, visualized in top views. The category with 9 trains is the most represented, accounting for 83% (n=29). **F**-**H)** Additional cryo-ET slices of undocked centrioles featuring attached ciliary vesicles and IFT trains leading to the ciliary vesicle membrane. **G)** Putative ciliary necklace particles are observed on the membrane of the ciliary vesicle (cyan arrowhead). **I**) Close-up view of putative ciliary necklace particles embedded within the ciliary vesicle membrane in (**G**)**. J**) Close-up view of individual IFT train connecting to the distal end of the undocked centriole and ciliary vesicle in (**G**). **K**) Template matching results (using an assembling IFT-B structure as initial reference; EMD-15258^35^) corresponding to the close-up view shown in (**J**) shows strong periodic correlation scores for the IFT train at the undocked centriole and ciliary vesicle. **L**) Subtomogram average of mTEC IFT trains (38 Å resolution), showing similar extended structural features to published anterograde IFT structures (**Figure S10E**).

We next used U-ExM to examine IFT at the ciliary base. IFT88 staining revealed a 9-fold symmetric occupancy of filamentous IFT trains extending from the transition zone microtubules into the cytosol, similar in appearance to assembling IFT trains in *C. reinhardtii*^35^ (**Figures 6E & S11**). It is noteworthy that the observed occupancy of assembling trains attached to the transition zone was less in our cryo-ET data than observed with U-ExM. This may be due to physiological differences in our mTEC and hTEC samples, perhaps caused by the different cell dissociation and handling protocols prior to vitrification or chemical fixation.

In addition to our observations of IFT at the ciliary base, cryo-ET imaging captured several instances of undocked centrioles below the cell’s apical surface– two of which were capped on their distal end by a ciliary vesicle^69,70^ (**Figure 6F-H**). In both cases, IFT trains were attached to the distal end of the centriole, beneath the ciliary vesicle. This observation suggests that IFT trains are recruited to the microtubules prior to vesicular docking with the plasma membrane and ciliogenesis. Close-up views of these vesicle-associated trains, template matching, and STA reveal features that are consistent with anterograde IFT trains (**Figures 6J-L**, **S10**). This provides *in situ* evidence for IFT trains assembling on undocked centrioles, suggesting they are primed for immediate transport initiation upon ciliogenesis. The ciliary vesicles contained smaller internal vesicles full of dense granular material and were decorated with arrays of membrane densities, some of which are consistent with ciliary necklace proteins (**Figure 6I**).

### Filamentous actin and intermediate filaments scaffold the apical surface of MCCs

Cryo-ET of MCCs revealed that ciliary microtubules are supported by an extensive cytoskeletal network of both filamentous actin and intermediate filaments. Bundled and branched actin filaments were present beneath the apical membrane and adjacent to the transition zone (**Figure S2**), including instances where bundles appeared to contact the ciliary membrane (**Figure 7A**). STA of traced actin bundles revealed filaments with a ∼7 nm diameter consistent with actin filaments, arranged in arrays along the membrane (**Figure 7B,D**).

**Figure 7:**
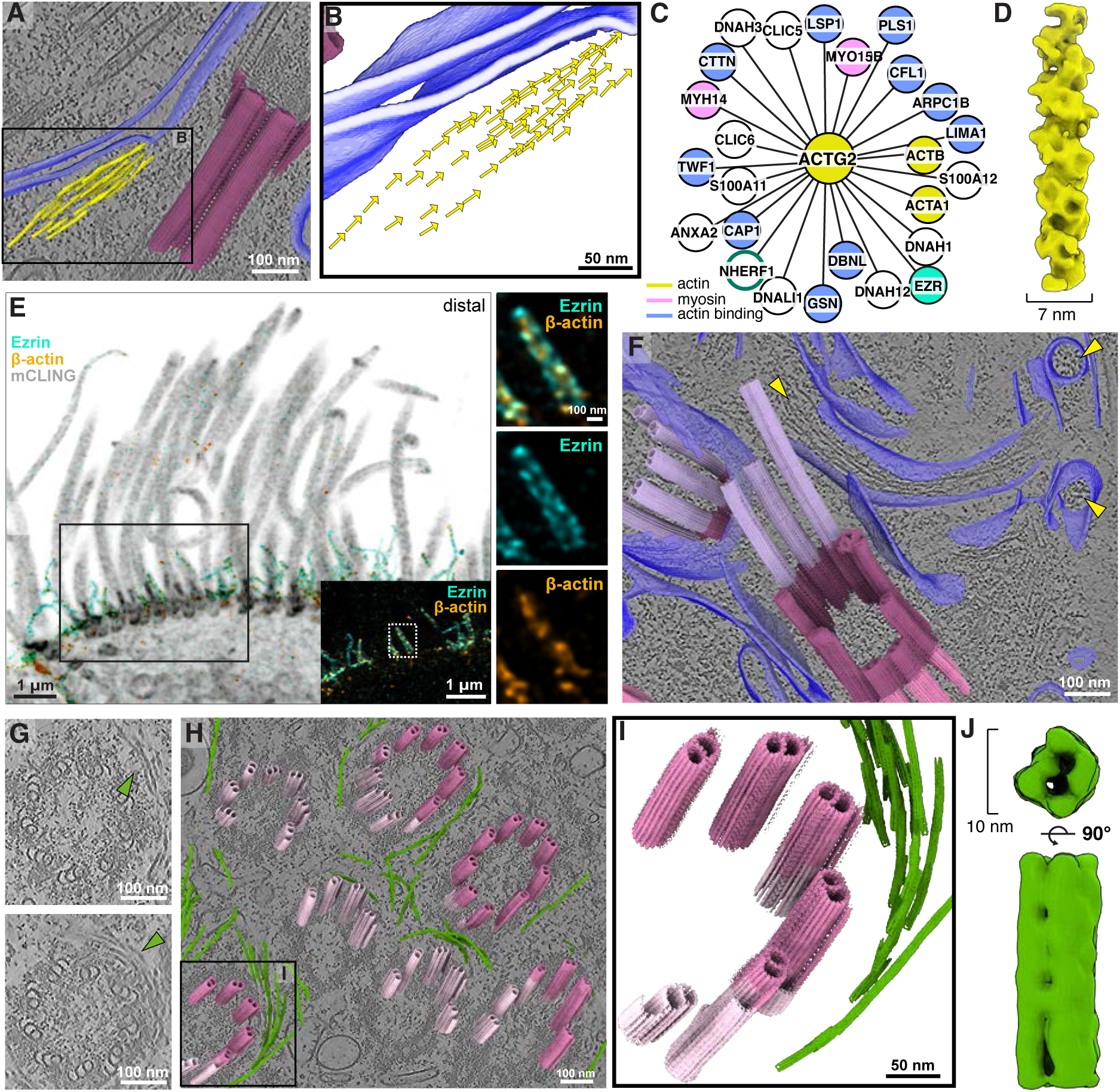
Actin and intermediate filament bundles organize the apical surface of MCCs. **A)** Cryo-ET of mTEC reveals bundled filamentous actin positioned under the apical membrane and at the ciliary pocket of MCCs. **B)** Mapbacks of averaged particle orientations is consistent with a parallel arrangement of actin filaments. **C)** Actin interaction network from XL/MS of bTEC shows association with several actin-binding proteins, myosin, and ciliary proteins. **D)** Subtomogram average of bundled actin (20 Å resolution, using helical symmetry) has a 7 nm diameter consistent with filamentous actin, but does not resolve polarity. **E)** Cilia of hTEC visualized by U-ExM and confocal microscopy. mCLING (grey) labels the membrane, Ezrin is in cyan, β-actin is in orange. **F)** Cryo-ET of mTEC shows cilia adjacent to microvilli structures. Ciliary microtubule mapbacks (shades of pink and purple as in Fig. 2) and membranes (blue) are superposed on the tomogram slice. Yellow arrows indicate filamentous actin in the microvilli. **G)** Cryo-ET identifies intermediate filaments (green arrowheads) wrapping around the ciliary base, level with the centriole. **H)** Mapbacks of centriole microtubules are shown with mapbacks of intermediate filaments. Inset: close-up around one centriole. **J)** Subtomogram average of the intermediate filament particle (27 Å resolution) reveals a hollow tube with a diameter of 10 nm, slightly wider than actin.

Our MCC interactome included a densely populated network of proteins that interact with actin. Analyzing proteins directly cross-linked to ACTG2 identified several actin-binding proteins, including PLS1, a protein in stereocilia that drives parallel actin bundling^71^ and may, by analogy, organize the membrane-proximal bundles observed in our cryo-ET data (**Figure 7C**). Although the STA resolution did not permit determination of filament polarity, previous work has shown that the barbed end of filamentous actin interacts with the membrane^72,73^.

U-ExM confirmed actin enrichment near the transition zone and at microvilli dispersed around the cilia, which were also observed in tomograms (**Figure 7E,F**). Co-localization with Ezrin revealed that the majority of actin is concentrated within the microvilli, with Ezrin lining the exterior (**Figure 7E**). We also detected Ezrin–NHERF1 cross-links, consistent with NHERF1’s known role in organizing membrane microdomains in MCCs^74^ (**Figure 7C**). Our data further revealed actin interactions with Ezrin, Moesin, and Radixin, which make up the ERM complex^75^ (**Figure 7C**) and are known to form links between the actin cytoskeleton and the plasma membrane, mirroring our observations from cryo-ET and U-ExM (**Figure 7A,E**).

In contrast, intermediate filaments were seen by cryo-ET to form a distinct layer further from the apical membrane, encircling the basal bodies at the level of the proximal and distal subregions (**Figure 7G-I**). STA resolved IFs as hollow filaments with a ∼10 nm diameter, readily distinguished from actin filaments (**Figure 7J**). Consistent with these images, our XL/MS analysis identified keratin IF proteins in proximity with ciliary proteins, in accord with previous localization studies of keratin isoforms to the MCC base^76^ (**Table S1**).

We further identified Annexin family proteins cross-linked to both actin and keratin intermediate filaments. Annexins have been implicated in ciliary membrane remodeling^77^, while keratins play roles in basal body spacing and cytoskeletal organization^78^. Together, our cryo-ET, U-ExM, and XL/MS data suggest that actin and IFs form a coordinated structural scaffold that may regulate ciliary spacing and mechanically integrate the ciliary base with the apical membrane.

## Discussion

By combining cryo-electron tomography, cross-linking mass spectrometry, and ultrastructure expansion microscopy, we built a molecular atlas of the ciliary base in mammalian MCCs (**Figure 8**). Some of structures we characterize were already apparent in the classical TEM images of ciliated epithelia from as early as the 1950’s ^20,70,79–81^. Our advance here is the ability to connect the morphology to molecular identity, interaction networks, and native three-dimensional organization. This is enabled by the complementarity of the three methods: cryo-ET revealed ultrastructure and spatial organization of macromolecular complexes, XL/MS identified proteins and their interaction partners with coarse spatial context, and U-ExM mapped the localization of individual proteins and their co-occurrence patterns. Taken together, these approaches allow us to move beyond description of individual structures and toward a molecular view of how the basal body, transition zone, ciliary membrane, and surrounding cytoskeleton are organized as a connected system. This builds on a recent shift in the field from isolated structural descriptions toward *in situ* analysis of ciliary base architecture and assembly.

**Figure 8:**
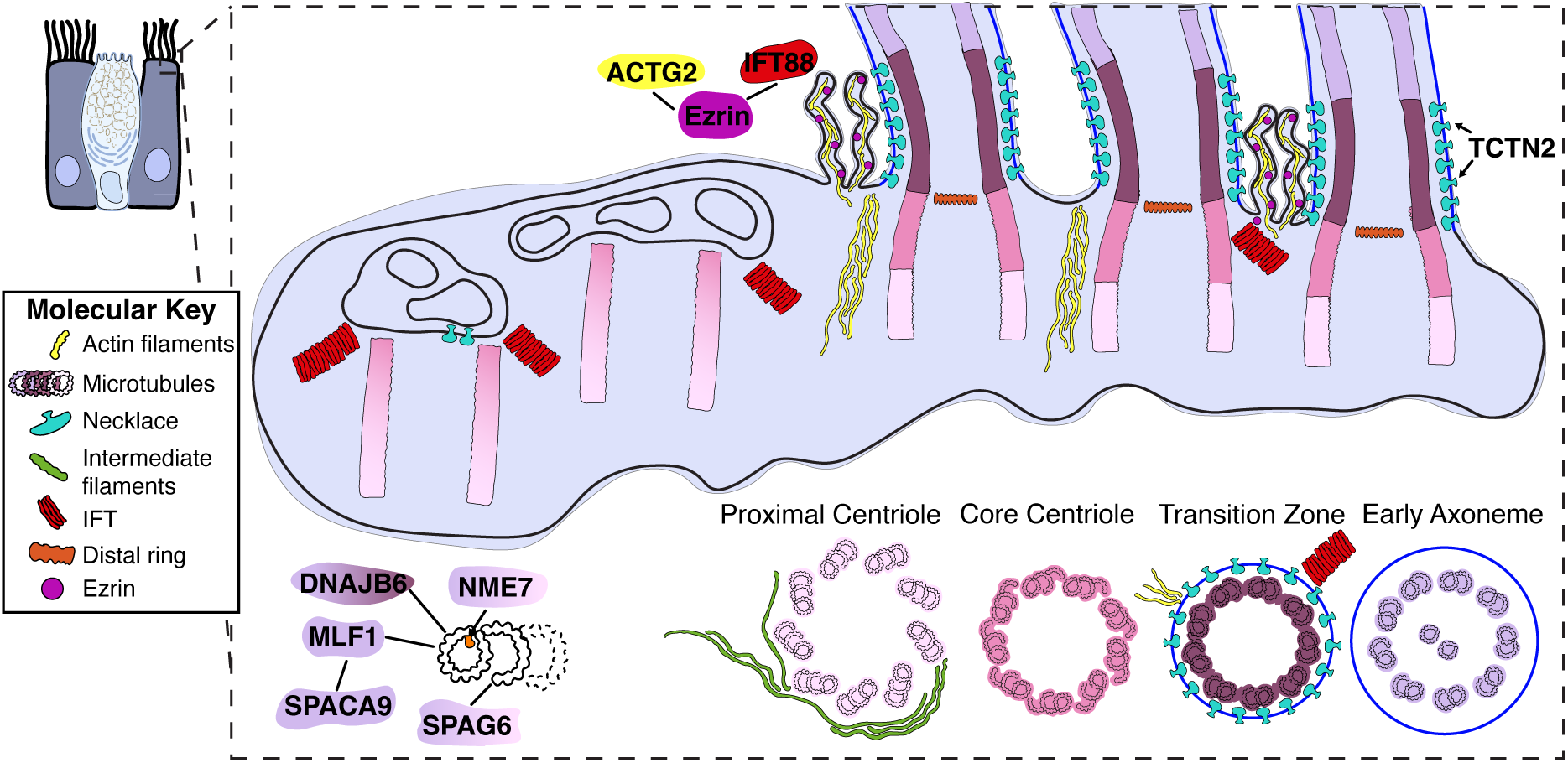
A molecular atlas of the MCC ciliary base architecture and interactions. The schematic summarizes the findings from this study including the structural changes of the microtubules along the ciliary ultrastructure, the ∼7 rows of the ciliary necklace adjacent to the transition zone, the presence of IFT trains at mature cilia and undocked centrioles, actin and intermediate filaments at the ciliary base, filamentous actin and ezrin in the microvilli surrounding the cilia, and several microtubule binding proteins.

Our analysis identified a set of tubulin-associated proteins not previously recognized as microtubule-associated in MCCs, and together they suggest a local proteostasis module at the proximal axoneme. MLF1^49,82^, which was previously noted in transcriptomic studies of ciliogenesis^51^, localizes to the ciliary axoneme and interacts with SPACA9, DNAJB6, and tubulin. SPACA9 is an established microtubule inner protein of both sperm singlet microtubules and the B-tubule of respiratory cilia^58^, suggesting that MLF1 may be recruited to a similar axonemal region. DNAJB6 is enriched at the transition zone and along the axoneme. Given its known role as an HSP70-associated anti-aggregation chaperone^83^, together with the linkage of SPACA9 to HSPA1B in our interactome, one possibility is that DNAJB6 participates in local quality control at the ciliary base, helping to stabilize or assemble proteins before their incorporation into the axoneme. This idea fits with earlier work on Hsp90α at the ciliary neck^84^, the TRiC chaperone tethered to the sperm axoneme^85,86^, and with the recently described role of DNAJB6 in quality control during nuclear pore biogenesis^87^. The different localization of DNAJB6 in primary cilia, which we report at the basal body, suggests that this chaperone may play a conserved but context-dependent role in ciliary assembly or maintenance. It remains an exciting future perspective whether MLF1 and SPACA9 work with DNAJB6 in this proteostasis capacity.

In addition to uncovering new MIPs and MAPs, our study provides molecular insight into the organization of the membrane at the ciliary base. Using cryo-ET, we visualized the ciliary necklace in MCCs as a regular array of membrane-associated structures closely aligned with the transition zone. Although Y-links are not sufficiently resolved in our averages to define their architecture directly, the close register of these two structures strongly suggests that they are functionally coupled. Quantification of the necklace indicates an ordered arrangement of approximately six membrane complexes per microtubule doublet, consistent with the idea that this membrane specialization contributes to the organization of the ciliary gate. This interpretation fits well with the long-standing view of the necklace as a specialized membrane domain^61,62^ at the transition zone and with more recent structural work in photoreceptors, where cryo-ET has resolved the necklace together with Y-links in the connecting cilium^32^. In that context, our localization of TCTN2 to the necklace region is notable, as TCTN-family proteins are established transition-zone components linked to membrane composition and gating^65,88,89^. It is also consistent with recent work identifying CEP290 in the membrane-proximal space external to the axonemal microtubules in the photoreceptor connecting cilium^32,90^, and showing that loss of CEP290 disrupts necklace organization^64^. Together, these observations support a model in which the ciliary necklace and transition zone form two closely apposed features of the same membrane–microtubule interface.

All three of our complementary approaches suggest a close relationship between IFT assembly and membrane organization at the ciliary base. The Ezrin-IFT88 cross-link is particularly informative. Ezrin is a membrane-actin linker^91^ whose apical localization in airway epithelial cells depends on FoxJ1^68^, and it contributes to organization of the apical membrane and to regulation of ciliary beating^74^. Its linkage to IFT88, together with our cryo-ET and U-ExM observations of IFT trains at the transition zone, suggests that anterograde IFT assembly occurs proximal to the ciliary membrane. This is consistent with recent structural work showing that IFT trains assemble stepwise at the ciliary base^35^. Importantly, we also observe IFT trains at the distal ends of undocked centrioles beneath ciliary vesicles, suggesting that recruitment of IFT machinery can begin before basal body docking. The necklace-like particles present on these vesicles further suggest that elements of the ciliary gate are preassembled on vesicular intermediates. Together, these observations support a model in which IFT assembly, membrane specialization, and gate formation are coordinated early in ciliogenesis.

We observe bundles of actin filaments at the ciliary base and in microvilli surrounding the cilia, and our XL/MS data identify a link between Ezrin and ACTG2, supporting the close association of actin with the apical ciliary membrane in airway MCCs. This is consistent with the distinct role of actin in MCCs^92,93^. Whereas actin polymerization generally antagonizes primary ciliogenesis^94^, in MCCs polymerized apical actin is required for basal body migration, docking, and mechanical stabilization of the ciliary array^95–97^. XL/MS identified several keratin proteins associated with ciliary components, and cryo-ET revealed many IFs wrapped around basal bodies, suggesting that the ciliary base is embedded in an extensive apical scaffold that includes keratin-based intermediate filaments as well as actin. This interpretation is consistent with earlier work showing extensive crosstalk between intermediate filaments and microtubules, including reorganization of keratin networks after microtubule perturbation^78^. In MCCs, such a composite scaffold would be well suited to anchor basal bodies and resist the mechanical forces generated by coordinated ciliary beating in a high-viscosity mucous environment.

Collectively, these data establish architectural principles for how MCCs achieve the organization required to support the assembly, anchoring, and coordinated beating of dozens to hundreds of motile cilia (**Figure 8**). This work provides a native view of the mammalian MCC base, while opening new biological questions. What transition zone-specific proteins stabilize features such as the A-B linker and the MIP helix? What roles do chaperones such as DNAJB6 play in ciliary assembly, function, and maintenance? How do Ezrin and the surrounding actin- and intermediate filament-rich cytoskeleton influence IFT dynamics, basal body docking, and mechanical stability? And to what extent are the features defined here conserved across motile and primary cilia, or instead adapted to the particular demands of the multiciliated state? More broadly, this work illustrates the value of integrative structural cell biology^38^. By linking native ultrastructure to molecular identity and interaction networks, it becomes possible to move closer to a mechanistic account of how complex cellular interfaces are built. Thus, the work described here advances understanding of mammalian MCC biology, provides a general framework for dissecting the molecular architecture of other subcellular systems, and generates cryo-ET and XL/MS resources for the community that are publicly available on EMPIAR and PRIDE databases to guide future ciliary studies.

## Methods

### MTEC cell culture

mTEC cells were obtained from primary culture of tracheal epithelial cells from wild-type CD-1 or transgenic GFP-centrin2 mice (University of Minnesota, Minneapolis, MN USA)^98^. After CO_2_ asphyxiation, ∼10 mice (per ALI culture) were dissected; trachea were isolated, cut open lengthwise, and stored in a petri dish with ice-cold PBS solution. After two washes with ∼10 mL of ice-cold PBS and removal of extraneous tissue, trachea were then incubated with 20 mL of 1.5 mg/mL Pronase E solution (in DMEM:F12 AB/AM) and incubated undisturbed for 18h (overnight) at 4 °C to dissociate epithelial and basal cells from the tracheal husks. After incubation, the flasks were gently inverted 2-5 times in order to further dissociate the epithelial and basal cells. 2.0 mL of FBS was then added to a final concentration of 10%, quenching enzymatic activity. The trachea/cell suspension was then vigorously inverted 12 times, for full dissociation of the cells. The trachea were removed using a Pasteur pipet, washed twice with two more rounds of 5 mL of fresh Ham’s F-12/Pen-Strep medium with 10% FBS, and inverted vigorously for 12 times each time. After removing the now empty tracheal husks, the three cell suspensions were pooled together and centrifuged at 600 x *g* at 4 °C for 10 min. Cells were resuspended in 1.0 mL DNAse, and incubated on ice for 5 min. After spinning down at 300 x *g* for 10 min at 4 °C in a microcentrifuge, pellets were resuspended in 20 mL of 10% FBS in DMEM:F12 AB/AM, and incubated for 4h at 37 °C, 5% CO_2_ in a 10 cm ø Primaria plate, to which fibroblasts would adhere, while leaving epithelial and basal cells in suspension. After this step, the cell suspension in the Primaria plate was gently swirled to detach non-adhered cells, and carefully transferred into two 15 mL Falcon tubes. The cells were spun down at 400 x *g* at room temperature for 10 min, then resuspended in ∼6 mL CM+RA.Corning Transwell Air-to-Liquid-Interface (ALI) culturing filter supports with collagen coating were shortly washed with CM+RA, after which 1 mL of cell suspension (in CM+RA) was added to each 6-well filter support insert (or 250 µL to each 24-well insert) to seed the primary culture. To the bottom well, below each filter support, 1.5 mL of CM+RA medium was added (0.5 mL in case of 24-well plates).

The cells on filter supports were incubated undisturbed at 37 °C, 5% CO_2_ for 5-6 days. Every second day, the CM+RA medium was refreshed, taking care not to disturb the cells on the filter supports. ROCK inhibitor^99^ was added to all CM+RA until day 4 of culture, after which it was left out. At day 5-6, confluency and compacting/columnar appearance of the newly formed cell layer was checked using a light microscope. If the cell layer had sufficiently developed, the switch to ALI culture was made by removing the medium in the filter support to create the air-to-liquid interface, and the medium in the bottom well was changed to 1.0 mL of either Nu+RA or SF+RA. The medium in the bottom well was then replaced every second day, and the air interface of the cell layer carefully rinsed with ∼1.5 mL of PBS every time to get rid of mucus and contaminants.

Ranging from day 7 to day 30 of ALI culture, filters were prepared for enzymatic dissociation by rinsing each well twice with BM. Trypsin solution (1.25%) was then added (0.5 mL to bottom well, 0.25 mL to filter support) and enzymatic dissociation was performed overnight (18 h) at 37 °C, 5% CO_2_. To quench the reaction, 0.25 mL of FBS was added to the pooled trypsinized cell suspensions, resulting in 1.00 mL of cell suspension. Cells were counted using a hemacytometer, while simultaneously checking for active ciliary beating, then spun down at 600 x*g* for 10 min at room temperature. The cell pellet was resuspended in BM with 10% FBS, in a volume that corresponded to the desired cell concentration for blotting onto EM grids, ranging from 125 to 500 cells per µL.

### HTEC cell culture

MucilAir™ *in vitro* tissue models of the human upper airway epithelium, cultured at the air-liquid interface, were purchased from Epithelix®. Cells were derived from trachea epithelial cells of a healthy 22-year-old male donor (batch number: MD0858). Upon receipt, MucilAir™ inserts were maintained in 24-well plates containing 700 µl of MucilAir™ culture medium and incubated at 37°C and 5% CO_2_ until further processing (within two weeks). The medium was changed twice a week. A weekly apical wash was performed by adding 200 µl of MucilAir™ medium to the surface of each insert. After a 30 min incubation at 37 °C and 5% CO_2_, gentle aspiration and dispensing were carried out to detach accumulated mucus and the mucus-medium was discarded.

### MTEC grid preparation

Using a Leica EM Grid Plunger (Leica Microsystems), 4 μL of ∼125-500 cells per μL of cell culture was mixed with 1 μL of 10- or 15-nm gold fiducial beads suspended in BSA solution, then blotted onto carbon-coated 200-mesh copper EM grids (Quantifoil Micro Tools). Blotting was performed at 95% humidity and 30 °C only from the back of the grid; a tiny drop of cell medium was applied to the back of the grid for better contact of liquid to filter paper, and therefore more reproducible wicking of the medium. The grids were plunge-frozen into liquid ethane.

The grids were then stored in sealed plastic containers under liquid nitrogen, and air-shipped in a liquid nitrogen-based dry shipping container. Upon arrival, grids were mounted in an Autogrid support, for further cryo-ET processing. Cryo-FIB sample preparation was performed using either a Scios or Aquilos dual-beam FIB/SEM instrument (FEI, Thermo Fisher Scientific). The grids were first coated with an organometallic platinum layer using a gas injection system. Subsequently, after milling stress relief trenches to improve FIB milling stability, the cells were milled with a gallium ion beam to produce ∼70- to 200-nm-thick lamellae, exposing the cellular interior.

### MTEC cryo-ET imaging

EM grids with FIB-thinned cells were transferred to a 300-kV Titan Krios microscope (FEI, Thermo Fisher Scientific), equipped with a post-column energy filter (Gatan) and a K2 Summit direct detector camera (Gatan). Using SerialEM software^100^, tilt-series were acquired from −60° to +60°, with 2° increments using a dose-symmetrical tilt scheme^101^. Images were recorded in movie mode at 12 frames per second, with an object pixel size of 3.52 Å (magnification of ∼42,000x) and a defocus of −4 to −6 μm. The total accumulated dose for the tilt-series was kept below ∼100 e^-^/Å^2^. Each tomogram was acquired from a separate cell and therefore can be considered a biological replicate. Several different cell cultures and 7 imaging sessions were used to produce the dataset of 72 tomograms.

### MTEC cryo-ET processing and subtomogram averaging

TOMOMAN^102^ was used to streamline the reconstruction of the tilt series^103–106^. Dose-weighting^105^ of tilt images was applied using TOMOMAN and tilt series were aligned using patch tracking in IMOD^106^. Tomograms were reconstructed in IMOD and then denoised using cryo-CARE^107^ followed by IsoNet^108^. CTF-corrected tomograms were also calculated using IMOD following the ctf3d routine.

Microtubules were traced manually using IMOD and oversampled every 4 nm along a spline using a script from STOPGAP^109^, always in the proximal to distal direction, based on the denoised tomograms. Particles were oriented such as to have the Z axis oriented along the filament and the in-plane angle randomized around the filament axis. Initial subtomogram averaging at “bin4” (14.08 Å/px) was performed in STOPGAP using subvolumes extracted from the CTF-corrected tomograms. The 3D classification initialized from randomly-assigned classes converged to five classes highly correlated to four well defined regions along the cillium: the proximal centriole, the core centriole, the transition zone and the early axoneme. Since the class assignment was not perfect, however, we employed custom scripts to reassign particle classes by voting among nearest neighbors, based on the known spatial constraints. Particle lists were then cleaned using an 8 nm distance using a STOPGAP script. Particle lists from individual classes were then imported into RELION-4^110^ for further refinement, including local alignments, CTF refinement and Bayesian polishing, which were iterated until convergence. Since initial defocus estimations were not accurate (as validated by inspecting the Thon ring fits from CTFFIND-4), especially for the higher tilt images, the option --reset_to_common in relion_ctf_refine yielded substantially improved maps. The transition zone linker map was obtained after recentering particles from the TZ MTD average to the linker density followed by local alignments. Given the large overlap between neighboring particles along the filament, the half-sets were split at the midpoint of each filament, minimizing spurious correlations in the Fourier Shell Correlation (FSC) curves while ensuring the maximum possible angular and defocus coverage in each half-set.

Ciliary necklace particles were manually picked from denoised tomograms that were segmented using MemBrain^111^ and visualized as membranograms with MPicker^112^. STOPGAP was used for initial subtomogram averaging and the average was further refined in RELION-4 using the same approach described above for the microtubule particles. Actin and IF particles were traced in IMOD, using the same approach as the microtubule particles, and averaged in STOPGAP before refinement in RELION-4 as described above.

Visual inspection of denoised tomograms resulted in 15 tomograms with putative IFT densities. Manual tracing of these trains proved difficult due to their low abundance and short length. We then performed template matching with pytom-match-pick^113^ on these 15 CTF-corrected tomograms (bin4) using the IFT-B subtomogram average from Chlamydomonas as a template (EMD-15258)^35^ at an angular sampling of 5 degrees (**Figure S10**). By maximum-intensity projection of the 15 score maps along Z, 5 tomograms revealed clear cross-correlation peaks for IFT trains, subsequently confirmed by visual inspection of the denoised tomograms, resulting in 427 candidate particles extracted and imported into RELION-4 for subtomogram averaging. 3D refinement with local searches yielded a subtomogram average at 38 `Å` resolution, which showed pronounced differences in density compared to IFT-B from the Chlamydomonas template. A second round of template matching was then performed using the native mTEC average as a reference, resulting in 183 true positive particles from 5 tomograms imported into RELION-4, which were subsequently refined to yield a subtomogram average still at 38 `Å` resolution but with higher map quality. FSC^114^ curves for resolution estimation were calculated using a soft-edged body-shaped mask for each map and are summarized in **Figures S10 & S12.**

### BTEC cilia isolation and cross-linking

Bovine trachea were collected immediately after slaughter and placed on ice for 10-30 minutes. The remainder of the preparation was performed in a 4 degree cold room. Trachea were cleaned of additional tissue and rinsed with cold PBS until the PBS ran clear. bTEC cilia were subjected to four separate preps –cilia and basal body enrichment, basal body enrichment, membrane and matrix enrichment, and microtubule enrichment-- to enrich for different ciliary proteins. bTEC trachea basal body prep followed previous preps where the trachea was first rinsed with cilia extraction PIPES buffer, followed by cilia extraction buffer as in ^115^. Basal bodies were then cleaned up by successive rounds of centrifugation at 2,000 xg and 13,000 xg to remove debris. Once the sample pellet was mostly white 8 mM DSSO in DMSO was added to the sample and incubated at room temperature for 1 hr, before being quenched with Tris buffer for 30 mins^3,94^. For the other preps, the cilia were isolated from the trachea as described in ^47^. After multiple rounds of centrifugation, the sample was cross-linked using the same protocol as described for the basal bodies. After the cross-linking and quenching, one third of the sample was aliquots for MS prep. With the remaining sample and additional subfraction step was performed as in ^116^ to enrich for either the membrane and matrix or microtubules.

### BTEC mass spectrometry

Enriched bTEC axonemes were resuspended in lysis buffer (5% SDS, 10mM TCEP, 0.1 M TEAB) and lysed by sonication using a PIXUL Multi-Sample Sonicator (Active Motif) with Pulse set to 50, PRF to 1, Process Time to 10 min and Burst Rate to 20 Hz. Lysates were incubated for 10 min at 95°C, alkylated in 20 mM iodoacetamide for 30 min at 25°C and proteins digested using S-Trap™ micro spin columns (Protifi) according to the manufacturer’s instructions. Shortly, 12 % phosphoric acid was added to each sample (final concentration of phosphoric acid 1.2%) followed by the addition of S-trap buffer (90% methanol, 100 mM TEAB pH 7.1) at a ratio of 6:1. Samples were mixed by vortexing and loaded onto S-trap columns by centrifugation at 4000 g for 1 min followed by three washes with S-trap buffer. Digestion buffer (50 mM TEAB pH 8.0) containing sequencing-grade modified trypsin (1/25, w/w; Promega, Madison, Wisconsin) was added to the S-trap column and incubate for 1h at 47 °C. Peptides were eluted by the consecutive addition and collection by centrifugation at 4000 g for 1 min of 40 ul digestion buffer, 40 ul of 0.2% formic acid and finally 35 ul 50% acetonitrile, 0.2% formic acid. Samples were dried under vacuum and stored at -20 °C until further use.

The sample was then enriched for cross-linked peptides using a GE Superdex 30 Increase 3.2/300 size-exclusion column (Cytiva) column. The dried peptides were resuspended in 30% acetonitrile, 0.1% TFA and the first 12 fractions were collected and dried for MS. Dried peptides were resuspended in 0.1% aqueous formic acid and subjected to LC–MS/MS analysis using an Orbitrap Eclipse Tribrid Mass Spectrometer fitted with a Ultimate 3000 nano system and a FAIMS Pro interface (all Thermo Fisher Scientific) and a custom-made column heater set to 60°C. Peptides were resolved using a RP-HPLC column (75 um × 30 cm) packed in-house with C18 resin (ReproSil-Pur C18–AQ, 1.9 μm resin; Dr. Maisch GmbH) at a flow rate of 0.3 ul/min. The following gradient was used for peptide separation: from 5% B to 13% B over 10 min to 38% B over 110 min to 95% B over 2 min followed by 18 min at 95% B then back to 5% B over 2 min followed by 18 min at 5% B. Buffer A was 0.1% formic acid in water and buffer B was 80% acetonitrile, 0.1% formic acid in water.

The mass spectrometer was operated in DDA mode with a cycle time of 4 s. Throughout each acquisition cycle, the FAIMS Pro interface switched between CVs of −40 V and −60 V with cycle times of 2 s and 2 s, respectively. MS1 spectra were acquired in the Orbitrap in profile mode at a resolution of 120,000 and a scan range of 375 to 1600 m/z, AGC target set to “Standard” and maximum injection time set to 50 ms. Precursors were filtered with monoisotopic peak determination set to “Peptide”, charge state set to 4 to 8, a dynamic exclusion of 30 s and an intensity threshold of 2e4. Precursors selected for MS2 analysis were isolated in the quadrupole with a 1.6 m/z isolation window and collected for a maximum injection time of 118 ms with normalized AGC target set to 200%, mass range set to “Normal” and scan range mode set to “Auto”. Fragmentation was performed with stepped HCD collision energies of 19%, 25% and 30% and MS2 spectra were acquired in the Orbitrap at a resolution of 60,000 in profile mode.

Cross-link search was performed using Scout^117^ against the corresponding database for each of the samples using the default values for DSSO-KSTY search with a CSM FDR cutoff of 1%. The search was repeated with xiSearch^118^ using the same database for DSSO-KSTY with a MS1 precursor tolerance of 6 ppm and fragment tolerance of 20 ppm and an CSM FDR cutoff of 5%. The consensus of these two searches was then used to build the final interactome.

#### HTEC expansion microscopy

Human airway epithelial tissues (MucilAir™, Epithelix) were used for all U-ExM experiments^41^. Prior to expansion, apical mucus was washed away using PBS. One MucilAir™ insert was placed into a 24-well plate containing 700 µl of Dispase II (5 U/ml in PBS1X) and incubated for 15 min at 37 °C and 5% CO_2_, to facilitate detachment of the tissue from the filter. The tissue was then removed from the filter by gently pushing it off with a razor blade and transferred into a 35 mm glass-bottom MatTek Petri dish, within the 10 mm microwell.

The tissue was incubated overnight at 37°C in 1 ml of 2% acrylamide + 1.4% formaldehyde in PBS1X. The solution was replaced with 90 µl of monomer solution (19% sodium acrylate, 0.1% bis-acrylamide, and 10% acrylamide in PBS10X) and incubated for 30 min at room temperature. The solution was then replaced with 90 µl of monomer solution supplemented with 0.5% TEMED and 0.5% APS. A 24 mm coverslip was placed over the microwell to seal it and trigger gelation, which was carried out for 15 min on ice, followed by 1.5 h at 37 °C.

After polymerization, the coverslip was removed, and the gel was trimmed around the tissue using a scalpel. The MatTek dish was then filled with 1 ml of denaturation buffer (200 mM SDS, 200 mM NaCl, 50 mM Tris H_2_O, pH 9) and incubated for 15 min at room temperature, until the gel detaches from the MatTek surface. The piece of gel containing the tissue was transferred into a 1.5 ml tube containing 1.5 ml of denaturation buffer, and incubated at 95 °C for 1.5 h. The gel was then washed in three 30 min baths of ddH_2_O and manually sliced with a razor blade to obtain approximately 0.5 mm thick transverse sections. Gel sections were incubated in two successive PBS1X baths before proceeding with immunostaining.

Gel sections were incubated overnight at room temperature under agitation with primary antibodies diluted in PBS1X containing 2% bovine serum albumin (BSA). Following four 15 min washes in PBS with 0.1% Tween 20, sections were incubated with secondary antibodies diluted in PBS-BSA for 2.5 h at 37 °C under agitation. A final round of four 15 min washes in PBS-Tween was performed, after which gel sections were expanded through three 10 min baths of ddH_2_O prior imaging.

Gel sections were mounted onto 24 mm coverslips coated with poly-D-lysine (0.1 mg/ml) and imaged with an inverted confocal Leica Stellaris 8 microscope using a 63× (1.4 NA) oil objective with Lightning deconvolution. Image processing was performed using ImageJ^119^.

The expansion factor (EF) was determined by measuring the width of 30 axonemes (5 from each of 6 different datasets), based on the tubulin signal, and comparing it to a reference value (i.e., 250 nm) using the PickCentrioleDim plugin. The resulting mean EF value (4.422) was then used to generate scale bars corrected for the expansion factor (**Figure S13**).

The average lengths of basal bodies and transition zones were measured using the PickCentrioleDim plugin. For BBs, the length was determined by measuring the GT335 signal from the proximal to the distal end of the basal body. The TZ length was calculated based on the shift between the distal end of the basal body and reappearance of the GT335 fluorescence at the proximal start of the axoneme. Based on n = 13 z-projections from N = 2 datasets (**Figure S14**)

### Data and Code Availability

Raw and reconstructed cryo-ET data for mouse tracheal epithelial cells is deposited at the Electron Microscopy Public Image Archive (EMPIAR) EMPIAR-13055. Subtomogram averages are available on the Electron Microscopy Data Bank (EMDB) XXXXX. Proteomics data is available on ProteomeXchange/PRIDE and MassIVE: MSV000101269.

## Acknowledgments

Calculations were performed at the sciCORE (http://scicore.unibas.ch/) scientific computing center at the University of Basel, Switzerland. We thank Jürgen Plitzko and Wolfgang Baumeister for access to microscopes and computing resources at the Max Planck Institute of Biochemistry, Martinsried, Germany. We thank the Biozentrum at University of Basel Proteomics Facility for support on methods development and data acquisition. This work was supported by SNSF Postdoctoral Fellowship TMPFP3_224900 awarded to C.L.M.; SNSF Project Grant PP00P3_187198 awarded to P.G. and V.H.; the Pro Visu and Gelbert Foundations supporting V.H.; ERC Consolidator Grant ISAC, administered in Switzerland by SERI (contract MB22.00075), awarded to P.G.; bridging postdoctoral funds provided by the Biozentrum to B.D.E.; and NIH grant R35GM130286 awarded to T.S.

## Author Contributions

T.S. and B.D.E. initiated cryo-ET experiments on mouse tracheal epithelial cells (mTECs). H.v.d.H. and G.B. prepared EM grids from mTEC cultures, then H.v.d.H. performed FIB milling and acquired the cryo-ET data, which was analyzed by C.L.M., A.M., P.V.d.S., R.D.R. and H.v.d.H. Ciliary isolations, biochemistry, and cross-linking on bTEC were performed by C.L.M. Mass spectrometry acquisition and methods development were implemented by C.L.M. and D.R. M.B. and O.M. optimized the U-ExM protocol for hTEC. M.B. performed all U-ExM experiments under the supervision of P.G. and V.H. C.L.M. wrote initial draft of the manuscript with comments and editing from all authors.

