## Supplement for "Molecular architecture of the ciliary base in mammalian multiciliated cells"

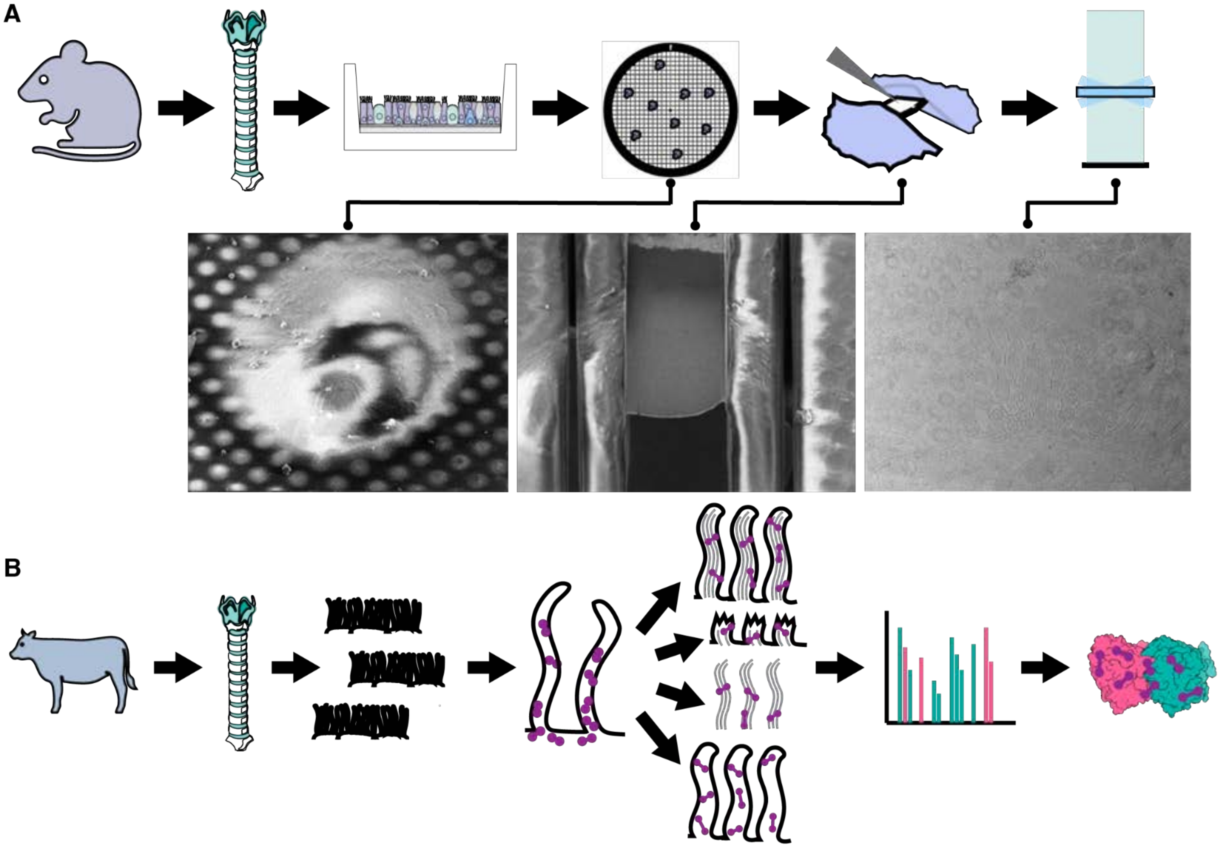


**Figure S1: Cryo-ET and XL/MS workflows. A)** Trachea were dissected from mice, then cells were dissociated from the tissue and grown on an air-to-liquid-interface (ALI). Enzymatic dissociation was used to isolate cells prior to plunge freezing. Plunge-frozen grids were thinned using FIB milling followed by tilt series acquisition on a 300-kV Titan Krios. **B)** Cow trachea were collected immediately after slaughter and all tissue was cleaned prior to ciliary isolation by mechanical shearing. Isolated cilia were cross-linked using DSSO before cilia were sub-fractionated into microtubule, membrane, and matrix fractions. Spectra were collected in DDA mode using FAIMS.

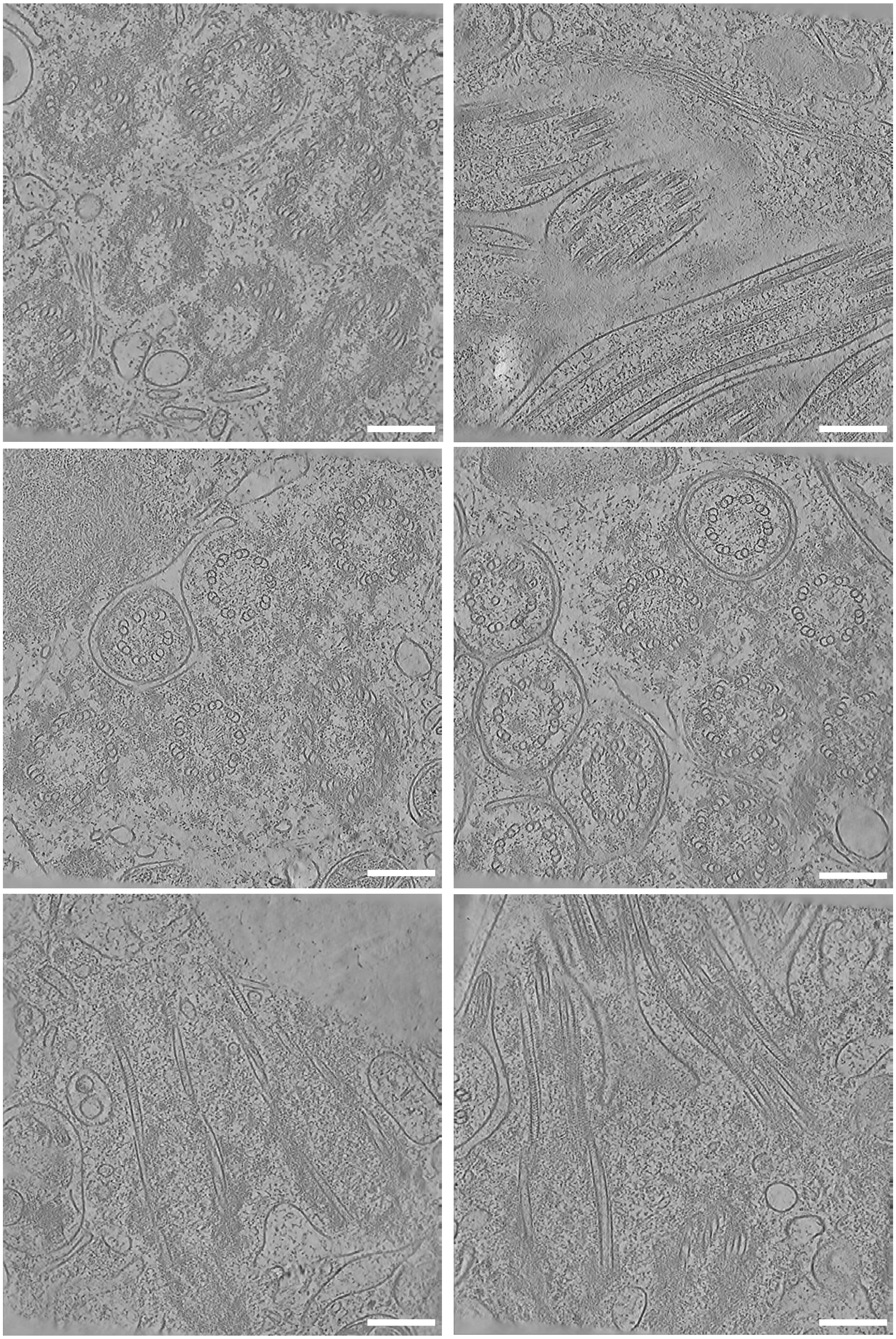

**Figure S2: Tomogram gallery of MTEC ciliary base.** Slices through selected tomograms, illustrating the complexity of the ciliary base. Scale bars: 200 nm.

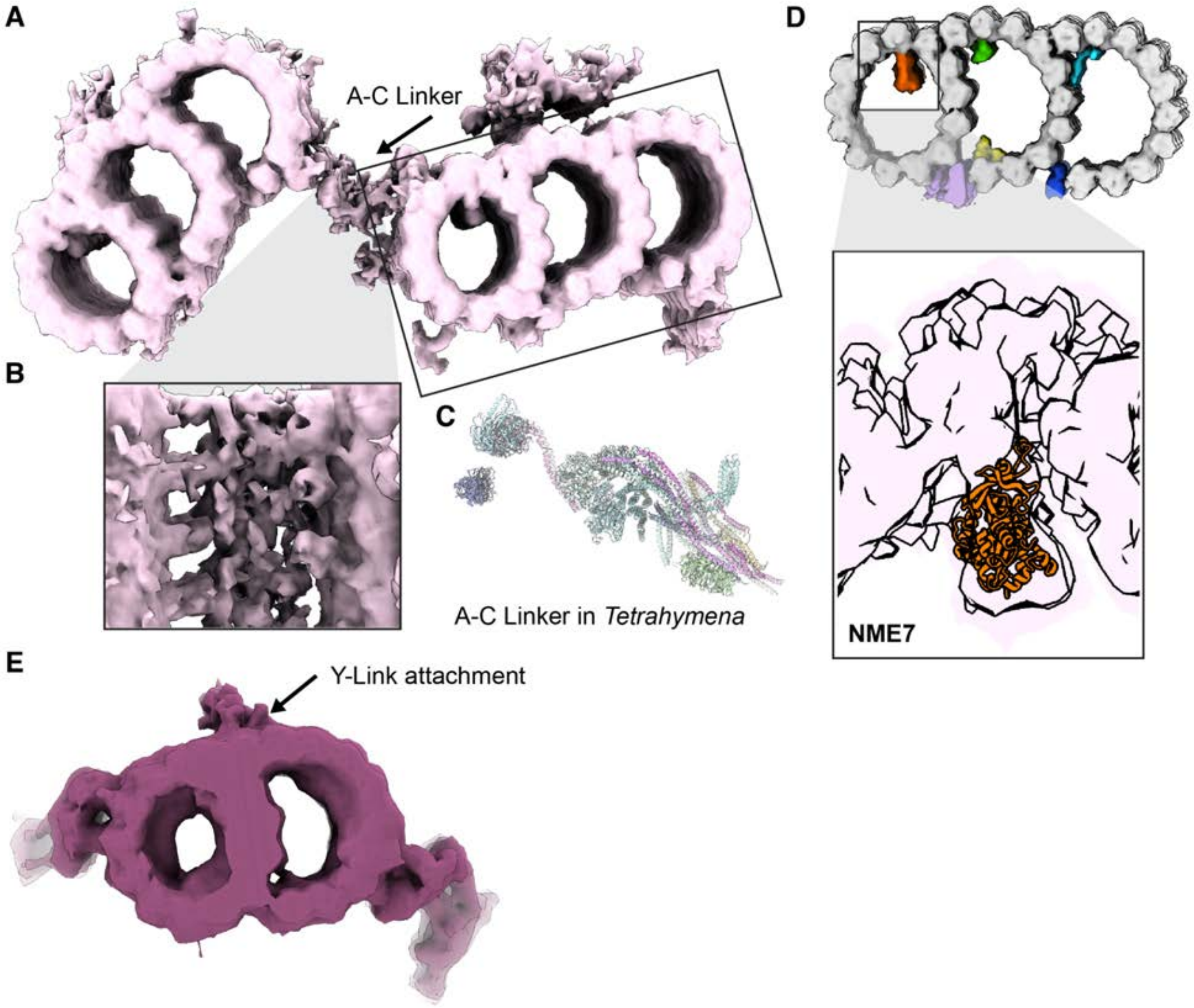


**Figure S3: Additional microtubule-associated densities. A)** The A-C linker from the STA map of the proximal centriole. **B)** Close up, side view of the A-C linker density. **C)** A-C linker structure from *Tetrahymena* (PDB: 9QZF), shown for comparison. **D)** Close up of the A9 MIP density in the proximal centriole with NME7 AlphaFold structure (orange) docked in. **E)** STA map of the transition zone, showing the connecting tip of the Y-link density.

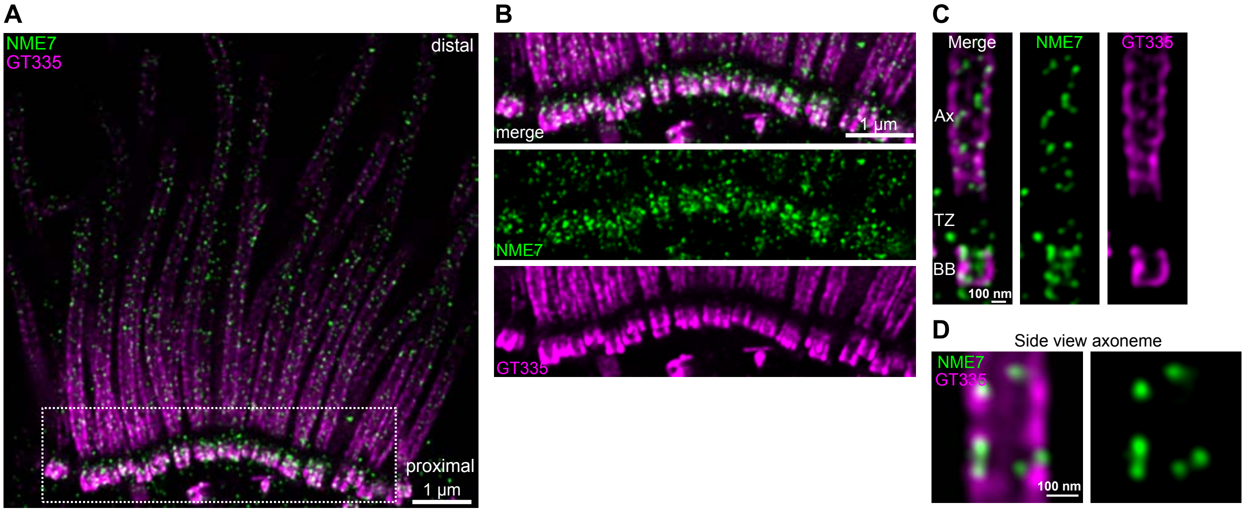


**Figure S4: U-ExM staining of NME7 in hTEC. A)** Overview of expanded hTEC, with NME7 in green and polyglutamylated tubulin (GT335) in magenta. **B)** Enlarged view of the region outlined in A. NME7 is highly enriched at the basal body and detectable along the axoneme but shows minimal presence at the transition zone (the gap in GT335 staining). **C)** Focus on the basal body (BB), transition zone (TZ), and base of the axoneme (Ax). **D)** Side-view of the axoneme.

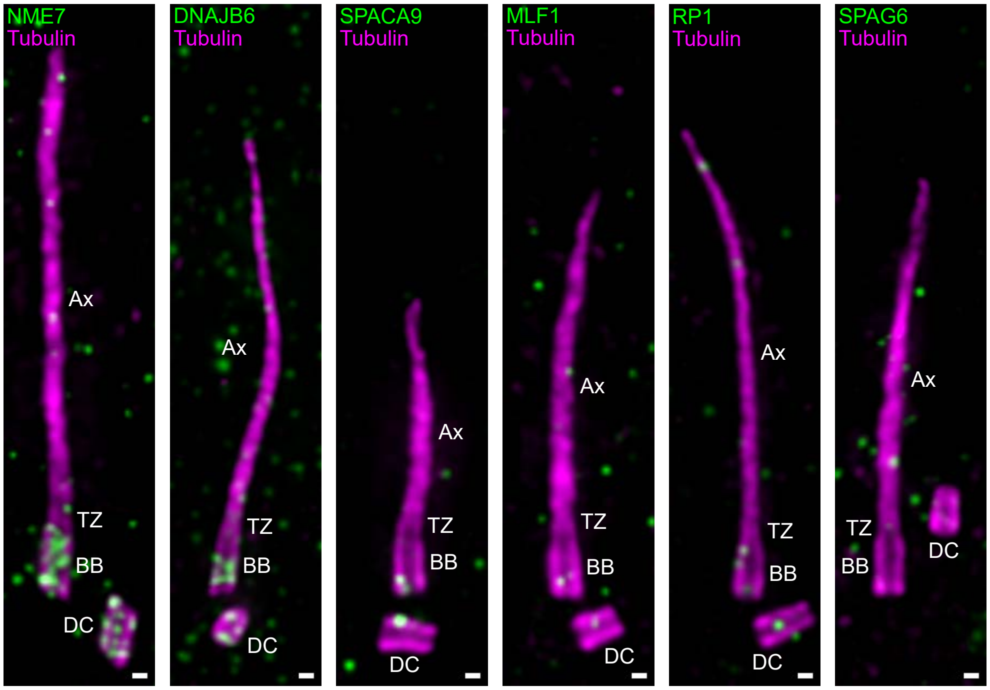


**Figure S5: U-ExM staining of NME7, DNAJB6, SPACA9, MLF1, RP1, and SPAG6 in hTERT RPE-1 cells.** Proteins of interest are shown in green and tubulin in magenta. NME7 and DNAJB6 both accumulate at the basal bodies (BB) and daughter centrioles (DC), whereas neither is detected at the transition zone (TZ) or along the axoneme (Ax). In contrast, SPACA9, MLF1, RP1, and SPAG6 show no specific signal in any ciliary sub-compartment. For each protein, n = 10. Scale bars are corrected for the expansion factor (EF = 4.8) and correspond to 100 nm.

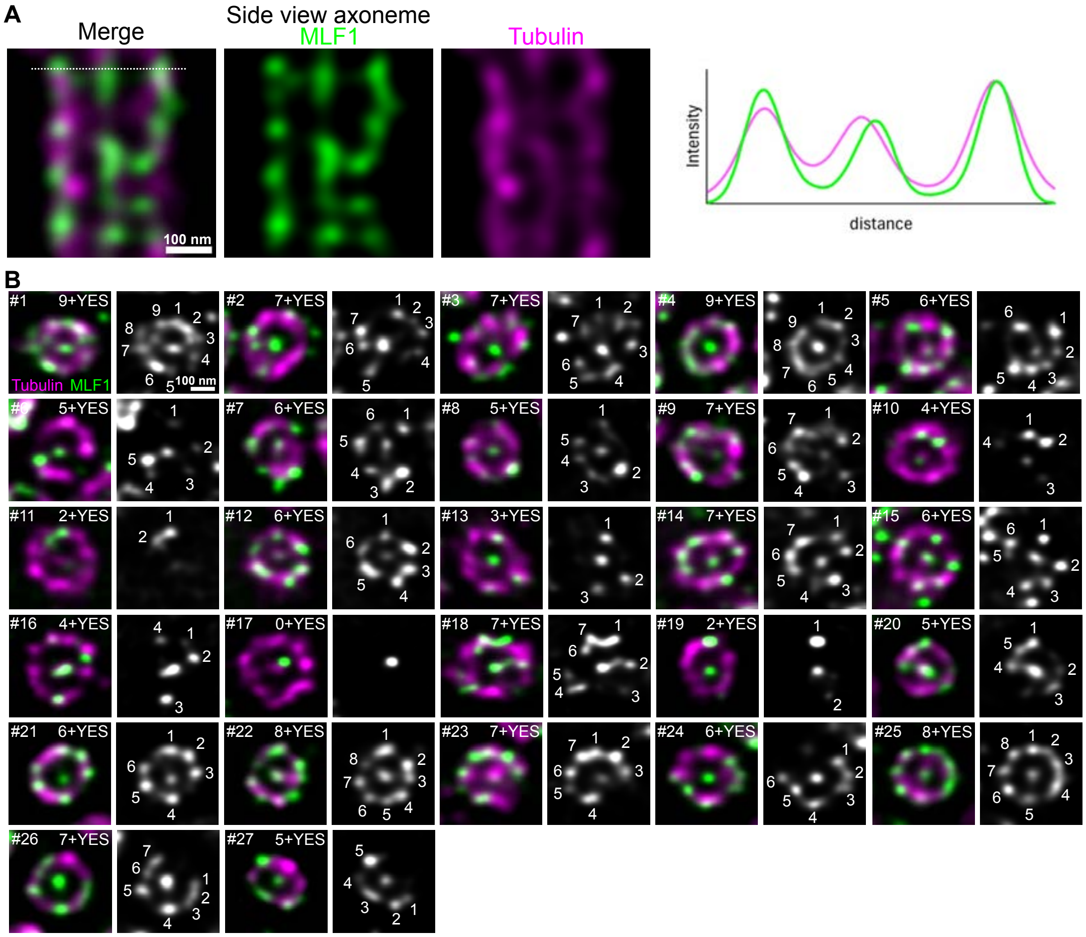


**Figure S6: U-ExM staining of MLF1 in hTEC. A)** Side-view of the axoneme (MLF1: green, tubulin: magenta). Intensity profile extracted from the region marked by the white dotted line. MLF1 is detected both along the microtubule wall and at the central pair. **B)** Top-view images of the axoneme used for the quantification showed in Fig 3 (greyscale: MLF1). The notation X+YES/NO indicates the number of MLF1 foci associated with the tubulin wall (X ranging from 0 to 9) and the presence (YES) or absence (NO) of MLF1 at the central pair. All analyzed axonemes show MLF1 signal at the central pair, with the most frequent category corresponding to seven foci associated with the tubulin wall (25%; n = 27.

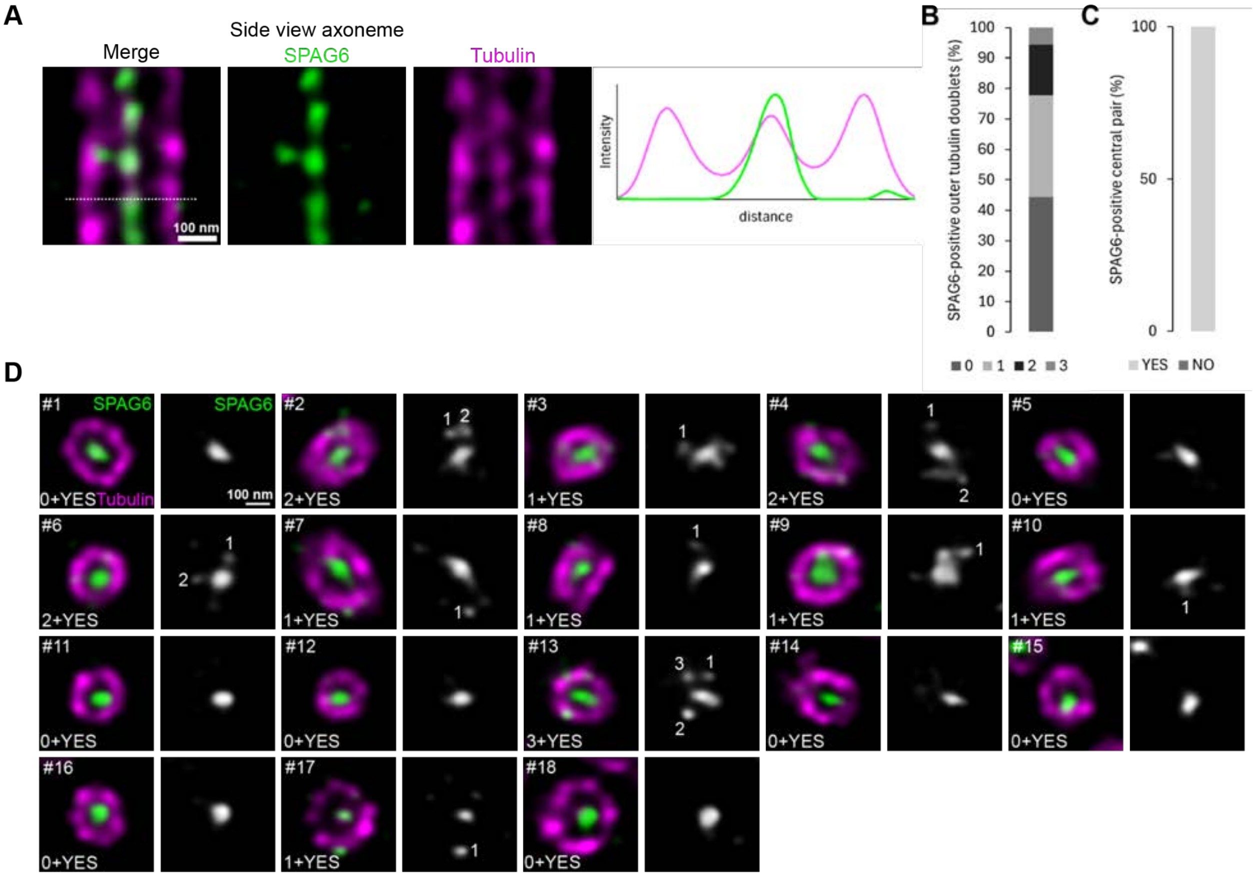


**Figure S7: U-ExM staining of SPAG6 in hTEC. A)** Side-view of the axoneme (SPAG6: green, tubulin: magenta). Intensity profile extracted from the region marked by the white dotted line. SPAG6 is detected at the central pair and occasionally at the outer doublets of microtubules that make up the axonemal wall. **B)** Quantification of SPAG6 foci associated with the tubulin wall, based on top-view images. In 44% of cases, SPAG6 is not detected at the outer doublets of microtubules; however, in some instances, 1 to 3 foci are observed (n = 18). **C)** Quantification of axonemes with SPA6-positive central pairs, based on top-view images. All analyzed axonemes show SPAG6 signal at the central pair (n = 18). **D)** Top-view images of the axoneme used for the quantification in B and C (greyscale: SPAG6). The notation X+YES/NO indicates the number of SPAG6 foci associated with the tubulin wall (X ranging from 0 to 9) and the presence (YES) or absence (NO) of SPAG6 at the central pair.

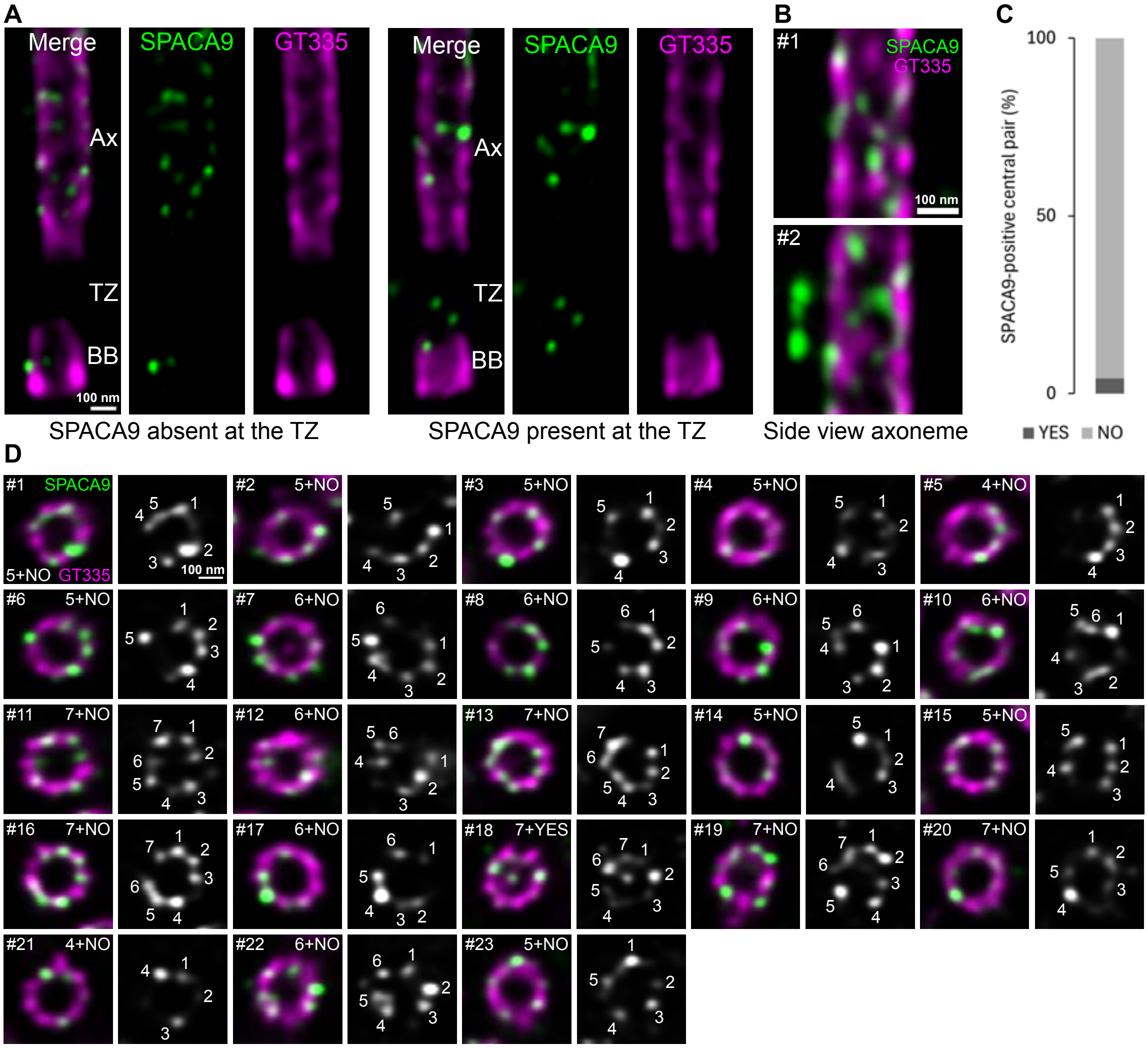


**Figure S8: U-ExM staining SPACA9 in hTEC. A)** Focus on the basal body (BB) - transition zone (TZ) – base of the axoneme (Ax). SPACA9 is detected at the axoneme for all cilia and for some at the BB and/or TZ. **B)** Side-view of the axoneme. In #2, an example showing SPACA9 accumulation along the axoneme, most likely at the level of the ciliary membrane, a recurrent localization. **C)** Quantification of axonemes with SPACA9-positive central pairs, based on top-view images. Among all the axonemes analyzed, only one showed SPACA9 at the central pair (n = 23). **D)** Top-view images of the axoneme used for the quantification in Fig 4F and C of this figure (greyscale: SPACA9). The notation X+YES/NO indicates the number of SPACA9 foci associated with the tubulin wall (X ranging from 0 to 9) and the presence (YES) or absence (NO) of SPACA9 at the central pair. In these images, the central pair of microtubules is not always visible because an antibody against polyglutamylated tubulin (GT335) was used instead of total tubulin, resulting in a less consistent labeling of the central apparatus. These top views were classified as axonemes based on their sufficient distance from the BBs.

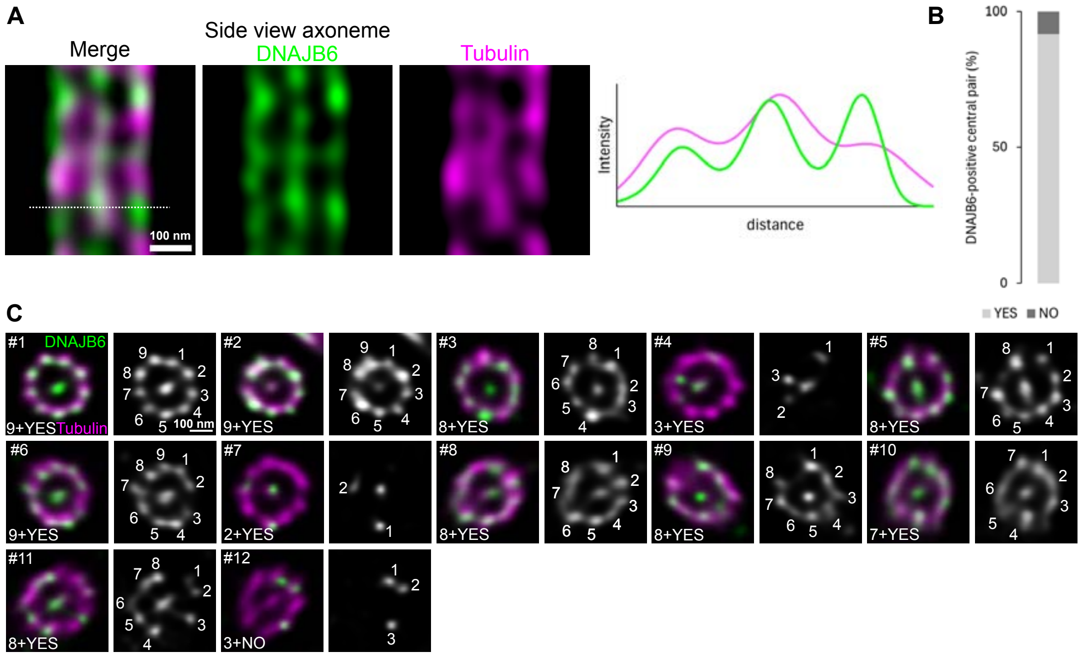


**Figure S9: U-ExM staining of DNAJB6 in hTEC. A)** Side-view of the axoneme (DNAJB6: green, tubulin: magenta). Intensity profile extracted from the region marked by the white dotted line. DNAJB6 is detected both along the microtubule wall and at the central pair. **B)** Quantification of axonemes with DNAJB6-positive central pairs, based on top-view images. DNAJB6 signal was detected at the central pair in 11 out of 12 axonemes examined. **C)** Top-view images of the axoneme used for the quantification in Fig 4H and B of this figure (greyscale: DNAJB6). The notation X+YES/NO indicates the number of DNAJB6 foci associated with the tubulin wall (X ranging from 0 to 9) and the presence (YES) or absence (NO) of DNAJB6 at the central pair.

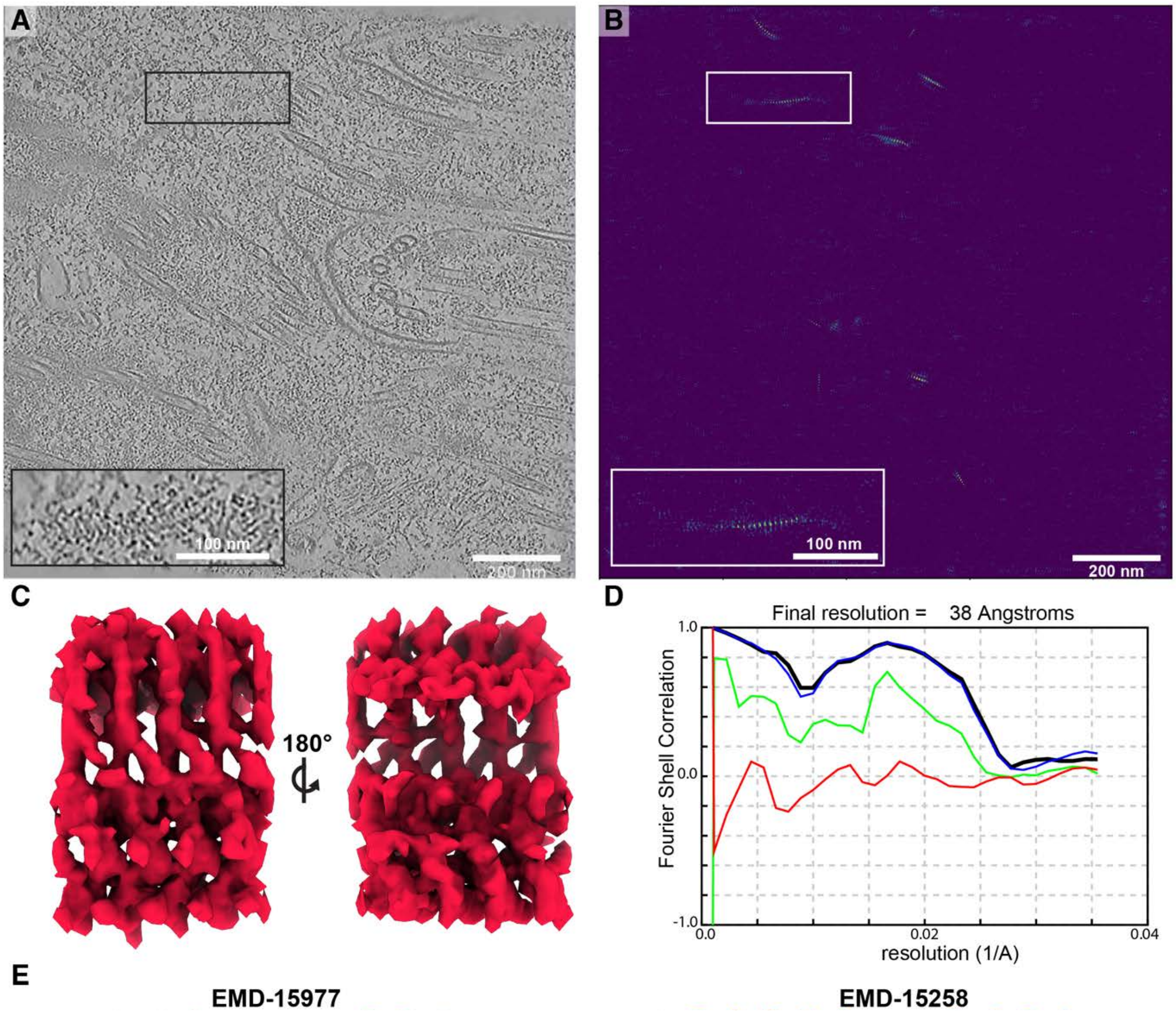


**Figure S10: IFT-B template matching and subtomogram averaging. A**) Representative tomogram with multiple IFT trains. Inset shows an IFT train identified with template matching. **B**) Score map from second round of template matching. IFT-B STA of assembling train from Chlamydomonas was initially used as a template (EMD-15258) before a second round of template matching was performed with template from the data and shows multiple periodic peaks in the regions where IFT trains are expected. The peaks corresponding to the train in (**A**) are shown in the inset. **C**) STA of IFT-B from this mTEC dataset, shown as an isosurface**. D**) FSC was calculated in RELION 4 at the 0.143 cutoff. Shown is the FSC corrected (black), FSC unmasked (green), FSC masked (blue), and FSC phase randomized masked (red). **E**) IFT-B averages from Chlamydomonas[35,67] highlight similarities and differences between the structures.

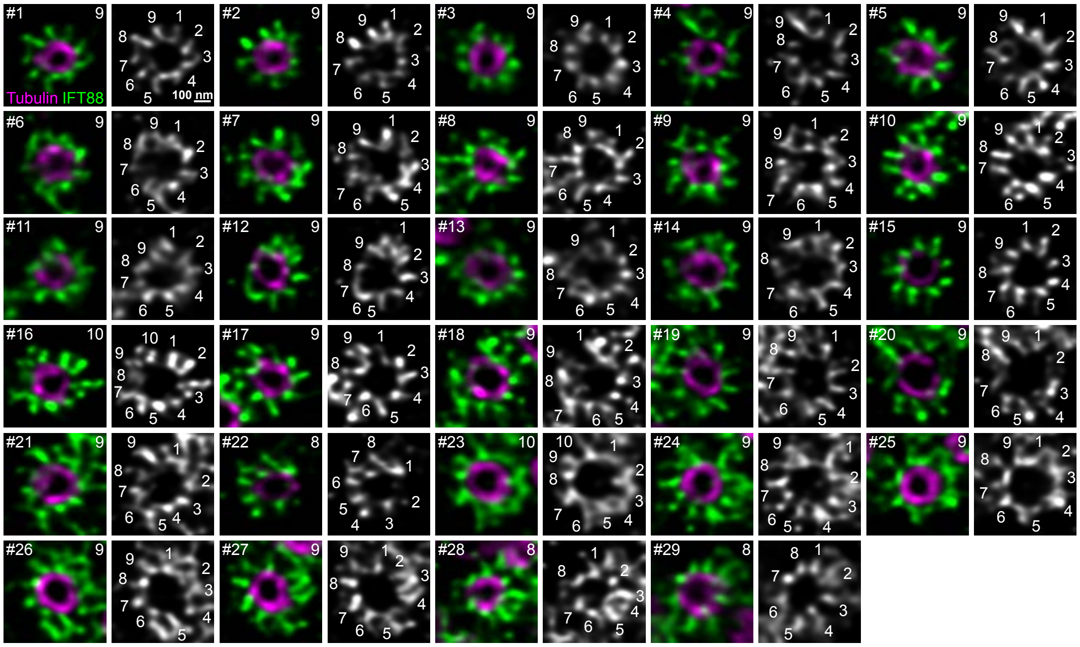


**Figure S11: U-ExM staining of IFT88 in hTEC.** Top-view images of the transition zone used for the quantification in Fig 6E (IFT88: green, tubulin: magenta; greyscale: IFT88). The notation indicates the number of IFT88 trains bound to tubulin (ranging from 8 to 10). The category with 9 trains is the most represented, accounting for 83% (n=29).

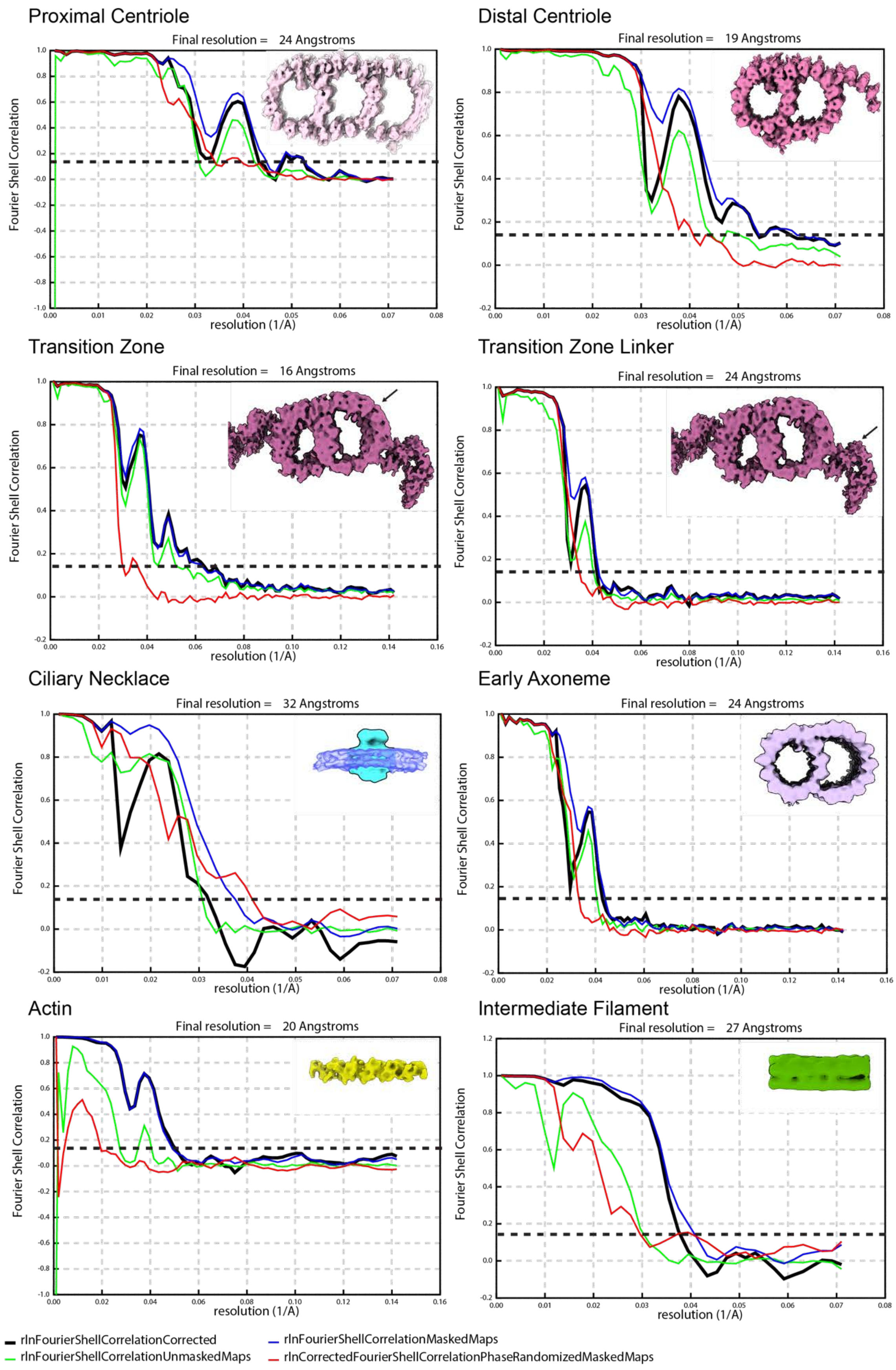
Proximal Centriole
Final resolution = 24 Angstroms
Distal Centriole
Final resolution = 19 Angstroms
Transition Zone
Final resolution = 16 Angstroms
Transition Zone Linker
Final resolution = 24 Angstroms
Ciliary Necklace
Final resolution = 32 Angstroms
Early Axoneme
Final resolution = 24 Angstroms
Actin
Final resolution = 20 Angstroms
Intermediate Filament
Final resolution = 27 Angstroms
Fourier Shell Correlation
resolution (1/A)
rlnFourierShellCorrelationCorrected
rlnFourierShellCorrelationUnmaskedMaps
rlnFourierShellCorrelationMaskedMaps
rlnCorrectedFourierShellCorrelationPhaseRandomizedMaskedMaps

**Figure S12: FSC plots for subtomogram averaging.** Fourier shell correlation is shown for each of the subtomogram averages from this work. FSCs were calculated in RELION 4 at the 0.143 cutoff. Shown is the FSC corrected (black), FSC unmasked (green), FSC masked (blue), and FSC phase randomized masked (red).

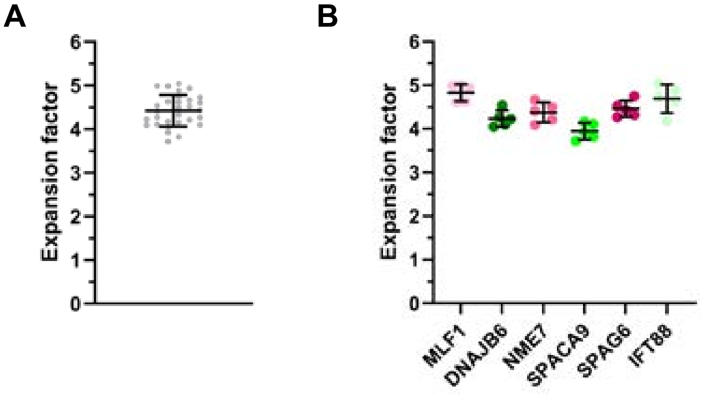


**Figure S13: U-ExM expansion factor for hTEC.** EF = 4.422 ± 0.3586 (mean ± standard deviation), based on n = 30 measurements from N = 6 datasets. **A)** All measurements combined. **B)** Each dataset is represented by a different color. MLF1 (light pink), NME7 (pink), SPAG6 (dark pink), IFT88 (light green), SPACA9 (green), DNAJB6 (dark green). **C)** Measurements grouped by dataset.

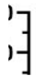

**Figure S14: Measurement of basal body and transition zone lengths in human tracheal motile cilia** based on the polyglutamylated tubulin (GT335) staining in U-ExM. BB: 268.9 ± 53.85 (mean ± standard deviation); TZ: 273.6 ± 54.73; BB and TZ: 542.5 ± 41.46 based on n = 13 measurements from N = 2 datasets.
